# *Escherichia coli* self-organizes developmental rosettes

**DOI:** 10.1101/2023.09.18.557913

**Authors:** Devina Puri, Kyle R. Allison

**Author notes:** Correspondence: Kyle R. Allison, **Email:**. Currently: Department of Molecular Microbiology and Center for Women’s Infectious Disease Research, Washington University School of Medicine, Saint Louis, MO. **Author Contributions:** DP performed experiments and analyses. DP & KRA designed experiments, interpreted results, and wrote the manuscript. KRA developed and performed motility experiments. **Competing Interest Statement:** The authors declare no competing interests.

## Abstract

Rosettes are self-organizing, circular multicellular communities that initiate developmental processes, like organogenesis and embryogenesis, in complex organisms. Their formation results from the active repositioning of adhered sister cells and is thought to distinguish multicellular organisms form unicellular ones. Though common in eukaryotes, this multicellular behavior has not been reported in bacteria. In this study, we discovered that *Escherichia coli* forms rosettes by active sister cell repositioning. After division, sister cells “fold” to actively align at the 2- and 4-cell stages of clonal division, thereby producing rosettes with characteristic quatrefoil configuration. Analysis revealed folding follows an angular random walk, comprised of ∼1-µm strokes and directional randomization. We further showed that this motion was produced by the flagellum, the extracellular tail whose rotation generates swimming motility. Rosette formation was found to require *de novo* flagella synthesis suggesting it must balance the opposing forces of *Ag43* adhesion and flagellar propulsion. We went on to show that proper rosette formation was developmentally required for subsequent morphogenesis of multicellular chains, *rpoS* expression, and formation of hydrostatic clonal-chain biofilms. Moreover, we discovered self-folding rosette-like communities in the standard motility assay, indicating this behavior may be general to hydrostatic environments. This study establishes that self-organization of developmental rosettes is a cross-kingdom multicellular behavior. Our findings indicate the potential of targeting bacterial rosettes to interrupt biofilms or reduce their antibiotic tolerance.

## Introduction

Multicellularity evolved from unicellular organisms billions of years ago, ultimately producing complex organisms like animals (1). “Multicellularity” is studied in diverse fields, but its meaning can vary from one to another (2). In a basic sense, an organism’s multicellularity results from its collected multicellular behaviors, which can range from simple (*e.g.*, cell-cell aggregation) to complicated (*e.g.*, organogenesis). Such behaviors are programmed (3), compared across species (4), experimentally evolved (5), and genetically dissected to understand multicellular development and evolution. Furthermore, identification of shared multicellular behaviors between dissimilar species could point to common design principles of multicellularity.

We recently discovered that *Escherichia coli* performs a facultative multicellular life cycle in hydrostatic environments (6, 7) (Fig. S1-2). Though *E. coli*’s relative multicellularity has long been considered (8, 9), our past study established several new multicellular behaviors (6). Tracking how an individual cell creates a multicellular biofilm, we observed the self-assembly of freely-moving multicellular chains (6). These chain communities were clonal, *i.e.* all cells descended form a single parental cell, and grew in length before later attaching to surfaces (6). We demonstrated this self-assembly had sequential morphogenetic stages, each regulated by a specific extracellular adhesin (6). Briefly, the self-recognizing outer-membrane protein Antigen-43 (*Ag43*) maintained sister-cell adhesion and clonality (starting at 2-cell stage); the extracellular polymer type-1 fimbriae controlled chain stability (∼8-cell stage); the amyloid-like curli controlled cellular positioning in chains (∼16-cell stage); and the extracellular polysaccharide poly-b-1,6-N-acetyl-D-glucosamine (PGA) attached chains to surfaces (∼1,000-cell stage) (6). Adapting a standard biofilm assay (7, 10), we showed hydrostatic *E. coli* biofilms resulted from this multicellular pathway (6), supporting the past idea that biofilms represent microbial parallels to *development* in multicellular organisms (11, 12). Several of *E. coli*’s newly-identified behaviors are common to complex organisms, which also develop through clonally-adhered division (4) and the stabilization of cellular positions by extracellular matrix (ECM) production (13). Additionally, *E. coli* cells were found to align in parallel to form square 4-cell “quatrefoil” communities (6), which we labeled “rosettes” following earlier descriptions of similar communities in mammalian host cells (14, 15). These rosettes partly enclosed an internal space and initiated multicellular morphogenesis (6), similar to developmental rosettes in multicellular organisms.

The rosette motif is ubiquitous in multicellular organisms and represents a key functional unit of tissue morphogenesis (16). Generally, rosettes are self-organizing, circular arrangements of cells that enclose an internal space and initiate developmental processes. They guide development across multiple scales (17), including of embryos (18), organs (19), and neurons (20–22). They are also thought to evolutionarily distinguish multicellular organisms from their unicellular relatives (4), and they are formed by many simple eukaryotes, including *Salpingoeca rosetta* (23), *Chlamydomonas* (24), and human parasites like *Toxoplasma* (25) and *Leishmania* (26). These examples are all flagellates that can perform clonally-adhered division, and *S. rosetta* in particular forms clonal rosettes by flagella-dependent repositioning of sister cells (27). Previously, it has been hypothesized that all animals descend from an ancient flagellate, capable of forming rosette-like communities, and that evolution of alternative repositioning mechanisms may have made the flagellum redundant (4). Hence, clonal rosette formation by cellular repositioning is a fundamental multicellular behavior performed by diverse eukaryotes. By contrast, this behavior has not been reported in bacteria. Though a variety of bacterial communities have been labeled “rosettes” (28–31), their formation is often unclear.

We considered the possibility that *E. coli* might initiate its multicellular life cycle by forming rosettes through cellular repositioning, similar to multicellular eukaryotes. Our past study showed that *Ag43* facilitated cell-cell adherence in rosettes while other adhesins, like fimbriae and curli, did not play a role until after rosette formation (6). However, *Ag43* adhesion alone should result in cells adhered solely at their poles (32) and is insufficient to explain quatrefoil configurations. This raises a potential role for *E. coli*’s flagellum, an extracellular tail that is rotated like a propeller by a molecular motor to produce swimming motility (33, 34). The flagellar motor switches between two rotational directions which separately cause swimming and tumbling, thereby producing a random walk (35, 36). The flagellum is required for hydrostatic biofilm formation, though its control by chemotaxis genes appears dispensable (37, 38). Despite this requirement, there was no evidence of swimming during the multicellular morphogenesis that generated hydrostatic biofilms (6), suggesting an alternative role for the flagellum. An example of flagella-assisted sister cell repositioning in *E. coli* was reported nearly 60 years ago (39) but, evidentally, was not further studied. Though our previous study reported *E. coli* rosettes (6), we did not rigorously investigate them given the incompatible time scales of their formation (minutes or less) and multicellular chain morphogenesis (hours). Here we combine microscopy, genetics, and quantitative analyses to investigate the mechanism and developmental consequences of *E. coli* rosette formation.

## Results

We used recently-developed methods (6) to record the motion of individual *E. coli* cells (model commensal strain K12 MG1655) by both bright-field and fluorescence microcopy. The resulting videos revealed *E. coli* rosettes form by a consistent behavior (Movies S1-2). Sister cells divided and adhered to each other at their new poles. After ∼10 minutes of negligible motion, cells “folded” onto one another, pivoting about their adhered poles to align in parallel (Fig. 1A) similar to previously-described motion (39). This folding often occurred in less than one minute and was comprised of angular movements interspersed with pauses. 2-cell “doublets” then grew and synchronously divided creating 4-cell groups, which folded while keeping their poles in contact to produce quatrefoil rosettes. No additional folding took place after 4-cell rosettes had formed (Fig. 1B). Instead, new sister cells adhere at their poles after division and transitioned into longitudinally growing chains while approximately keeping quatrefoil configuration. These findings demonstrate that rosettes are generated by sister-cell folding at the 2-cell and 4-cell stages. Cellular folding, combine with the stabilization of resulting quatrefoil structure by ECM production (6), provides an explanation for the well-defined morphogenesis of *E. coli* multicellular chains.

**Figure 1.**
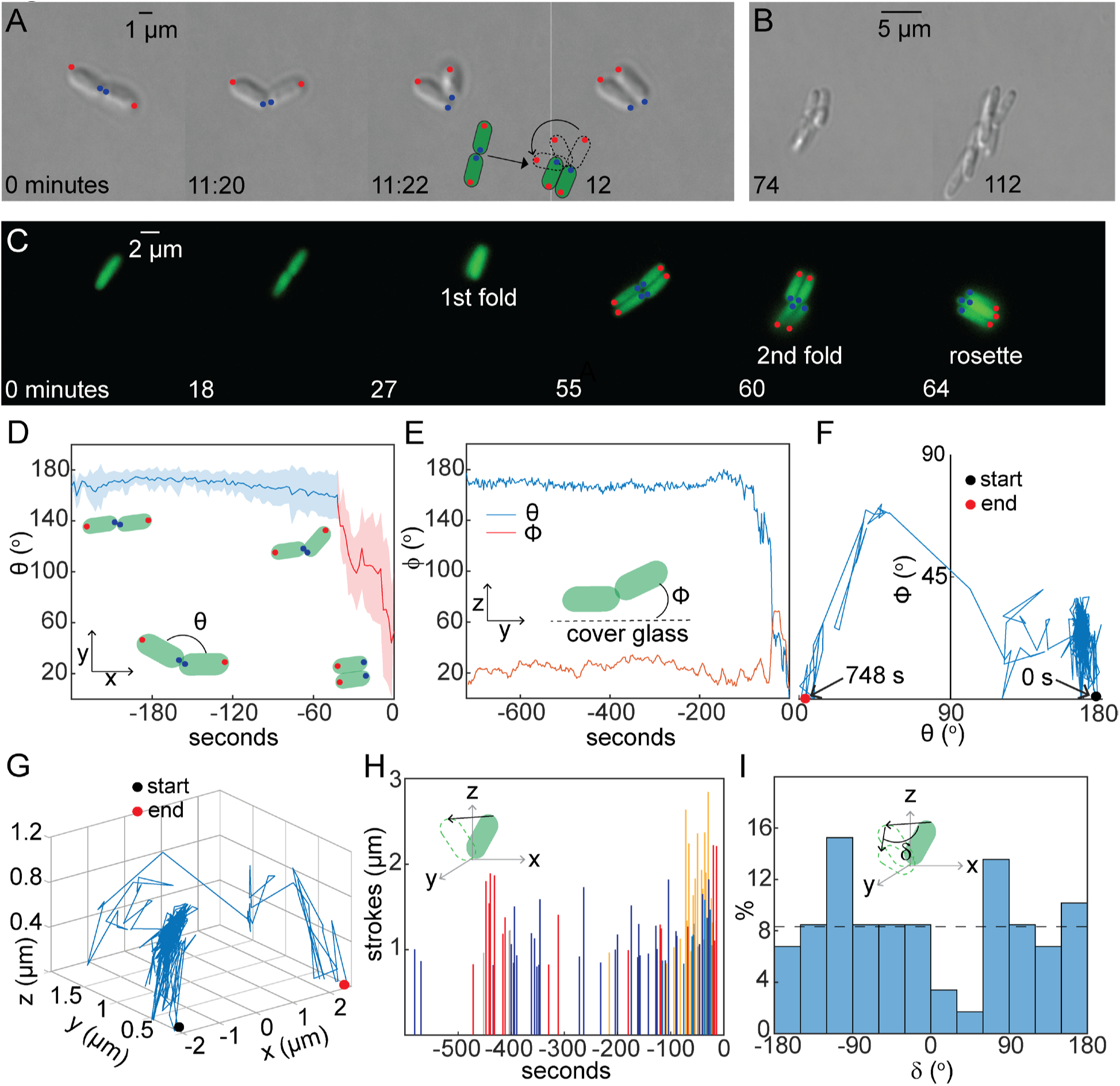
Sister-cell folding produces rosettes and follows an angular random walk. (A) Micrographs from a representative movie (2-second resolution) of *E. coli* sister-cell folding and parallel alignment. Time and scale indicated on all micrographs. New (blue) and old (red) poles are indicated. Inset: illustration of cell folding documented in movies. (B) Later micrographs from the same video. (C) Micrographs from a representative movie (30-second resolution) of rosette formation in cells constitutively-expressing GFP. New (blue) and old (red) poles are indicated. (D) Dynamics of sister-cell angle (θ) during cellular folding (2-second resolution; mean ± standard deviation, n=5). Folding (red) and pre-folding (blue) are indicated. Inset: illustration of θ calculation. (E) Representative example of sister cell θ (blue) and Ф (red). Inset: illustration of Ф calculation. (F) Trajectory of sister-cell angles from representative example of folding. Start and end positions and times are indicated. (G) Representative example of 3D trajectory of cell pole during folding. (H) Dynamics of angular strokes (movement >0.8 µm) for 4 representative sister-cell pairs. Inset: illustration of distance calculation. (I) Distribution of angle change between strokes (δ) (59 recorded strokes across 5 independent sister cell pairs). Inset: illustration of δ calculation.

To quantify folding, we next fluorescently imaged cells containing green fluorescent protein to enable image processing (GFP) (Fig. 1C; Fig. S3-8; Movies S3-7) and calculated the 2-dimensional angle between sister cells (θ) (Fig. S9). Changes in θ were small immediately after division, but greatly increased as sister cells folded into parallel orientation (Fig. S9 & 11). Circumferential rotation of cells around their long axis (“spinning”) (Movie S3) was also noted during pauses, further indicating folding is a combination of angular movements and paused spinning. We calculated the delay period between cell division and the onset of folding to be ∼13 minutes (Fig. S11). The maximum angular speed was >6°/s, the detection limit based on the 30-second imaging rate (Fig. S10-11). These findings demonstrate that sister-cell folding is comprised of two distinct motions and is regulated relative to the cell cycle, but also indicate that faster, three-dimensional data will be required.

To further quantify sister-cell folding, we recorded bright-field videos at 2-second resolution (Fig. S12-16; Movies S8-12), and tracked the position of cell poles. The angle between a cell and the cover glass (Ф) can be imputed by trigonometry from the cell’s length and 2D-projection. Together, θ and Ф values fully specify the 3-dimensional angle between sister cells. Calculated θ dynamics (Fig. 1D; S17) were consistent with the fluorescent data (Fig. S11), revealing delayed-onset sister-cell folding. The cell-averaged maximum angular speed of θ was calculated to be ∼36°/s (Fig. S18). Ф dynamics (Fig. S19) were similar to θ dynamics and significant changes in Ф and θ coincided (Fig. 1E; Fig. S20). The angular trajectories of sister cells (plotting Ф against θ) suggested folding may perform an angular random walk (Fig. 1F; Fig. S21). We calculated trajectories of the un-adhered cell poles in Cartesian coordinates, which also suggested random-walk motion (Fig. 1G; Fig. S22-23) and illustrated that folding resulted from switching between angular strokes and paused directional randomization. Inter-stroke pause times fit a long-tail distribution (Fig. S24) with similar time scales to rotational switching of the flagellum (40–42). Stroke distance was 0.8-1.2 µm (Fig. 1H), though in some cells increased to ∼2 µm immediately before folding completed. The angle between consecutive strokes (δ) was found to be uncorrelated (r=-0.041; n=59) and near uniformly distributed (Fig. 2I; Fig. S25) satisfying the criteria for a simple random walk (43).

**Figure 2.**
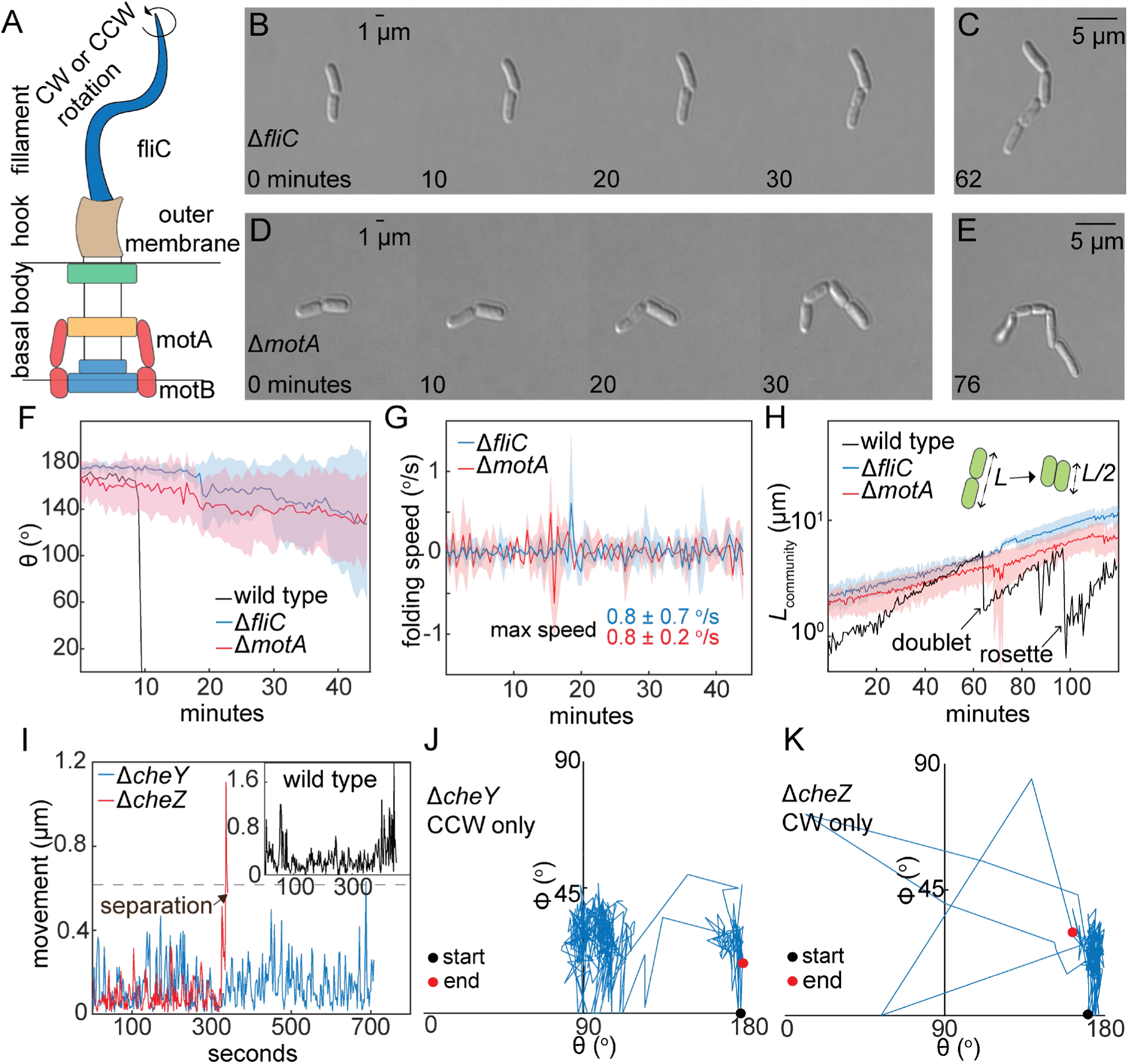
Flagella propulsion generates sister cell folding. (A) Diagram of the flagellum (not to scale). Micrographs from representative movies of (B) Δ*fliC* and (D) Δ*motA* cells. Time and scale indicated on all micrographs. Later micrographs of (C) Δ*fliC* and (E) Δ*motA* cells from the same videos. (F) Dynamics of Δ*fliC* (blue) and Δ*motA* (red) sister-cell angle (30-second resolution; mean ± standard deviation, n=3). An example wild-type trajectory is included for comparison. (G) Dynamics of angular speed for Δ*fliC* and Δ*motA* cells (30-second resolution; mean ± standard deviation, n=3). Inset: average maximum speed during. (H) Length dynamics of Δ*fliC* and Δ*motA* communities (mean ± standard deviation; n= 3). An example wild-type trajectory is included for comparison. Inset: illustration of halving length during folding. (I) Dynamics of angular movement for representative Δ*cheY* (blue) and Δ*cheZ* (red) sister-cell pairs (2-second resolution). Inset: representative wild-type example. (J) Trajectory of sister-cell angles from representative Δ*cheY* example. (K) Trajectory of sister-cell angles from representative Δ*cheZ* example.

Based on this motion and the considerations provided in the introduction, we hypothesized that sister-cell folding was generated by the flagellum (Fig. 2A), and we tracked non-motile Δ*fliC* and Δ*motA* strains (Movies S13-14). Δ*fliC* cells lack the main tail protein while Δ*motA* cells lack a motor component and are deficient in propulsion. Initially, Δ*fliC* and Δ*motA* cells behaved similarly to wild-type: they grew, divided, and adhered at their poles (Fig. 2B & D). However, neither Δ*fliC* nor Δ*motA* cells produced angular movements or folded into rosettes. Instead, Δ*fliC* and Δ*motA* cells grew in linear, unfolded chains where cells were solely adhered to each other at their poles (Fig. 2C & E). Quantifying θ demonstrated negligible angular motion in Δ*fliC* and Δ*motA* cells (Fig. 2F & G). Angular motion at the 2-cell and 4-cell stages in wild-type, and its lack in Δ*fliC* and Δ*motA*, was further demonstrated by quantifying community length, which suddenly halves when sister-cell folding occurs (Fig. 2H; Fig. S26). These findings demonstrate that *E. coli* sister cells reposition themselves into rosettes by flagella propulsion, similar to *S. rosetta* (27). Similar mechanisms in the bacterium *E. coli* and eukaryote *S. rosetta* indicate that clonal self-assembly of rosettes by flagella propulsion is a cross-kingdom multicellular behavior.

We next sought to elucidate the precise connection between flagella rotation and cellular motion during folding. The flagellum can rotate in both the clockwise (CW) and counterclockwise (CCW) directions, causing tumbling and linear swimming, respectively. We hypothesized that CW and CCW directions produced angular strokes and directional randomization, respectively. To test this, we used Δ*cheY* (mainly CCW) and Δ*cheZ* (mainly CW) mutants (44, 45). Videos demonstrated that the Δ*cheY* mutation abolished angular stokes (Fig. S26-30; Movies S15-17). However, some Δ*cheY* cells “pushed” past each other to align in parallel, though with an opposite polar orientation to folding. This push motion resembled the expected CCW-generated swimming of this strain (44, 45). Quantifying motion of Δ*cheY* sister cells revealed that travel distance remained <0.6 μm (Fig. 2I) and was comparable to the random motion of wild-type cells before they began folding (Fig. 2I, inset). Sister-cell angular trajectories in Δ*cheY* further demonstrated that angular strokes were abolished and illustrated that pushing rather than folding was responsible when parallel alignment did occur (Fig. 2J). We next tracked Δ*cheZ* sister cells and found that this mutation abolished the pauses between angular strokes (Fig. S31-34; Movies S18-20). Moreover, Δ*cheZ* sister cells often forcibly separated, similar to previously-described motion (39), thereby blocking community formation (Fig. S35). Quantifying sister-cell motion and angular trajectories further demonstrated that Δ*cheY* mutation abolished the pauses and caused cell separation (Fig. 2I & K). Together, these findings confirm that CW flagella rotation produces the angular strokes of sister cells while CCW flagella rotation produces their directional randomization. This reflects an inversion of the functional roles of flagella rotation in chemotaxis where CCW causes swimming (directional motion) and CW causes tumbling (directional randomization) (35, 36).

Though these findings reveal how flagella rotation produces the motions of sister-cell folding, the role of switching between rotational directions is unclear. CW rotation alone should produce sufficient motion to fully fold sister cells, and CCW should therefore be dispensable. However, when CCW rotation is absent, CW rotation forcibly separates sister cells and blocks rosette formation. Taken together, this may suggest switching itself serves a role in rosette formation, possibly by “braking” the angular motion. Such braking would prevent sister-cell separation and increase the probability of rosette formation in wild-type relative to Δ*cheZ* cells. To explore this, we quantified the frequency of community configurations after several generations (Fig. S36), and found that ∼83% of wild-type, ∼29% of Δ*cheZ,* ∼0% of Δ*cheY* communities were rosettes or rosette-configured chains. The remainder of communities were either single-cell-wide chains or aggregates lacking rosettes configuration. These results indicate that rotational switching itself is important to rosette folding, but not an absolute requirement. In the future, the precise role of motor switching in rosette formation could be determined by controllably varying its rate (40). Moreover, these findings are consistent with past data indicating differences between flagella and chemotaxis genes in hydrostatic biofilm formation (37, 38, 46, 47). It was recently suggested that *E. coli* combines *Ag43* adhesion and AI-2-based chemotaxis to form polyclonal aggregates (46). Our findings do not exclude this possibility, but instead demonstrate that rosettes form by a separate mechanism that also utilizes *Ag43* adhesion and flagellar propulsion.

The separation of sister cells by unchecked CW rotation (Δ*cheZ*) suggested flagella propulsion can overcome the adhesive forces holding cells together. Consistently, a role for flagella propulsion in sister cell separation has been described (39). These observations suggest that there may be an optimal amount of force enabling rosette formation. The force produced by a flagellum corresponds to its tail length and number motor subunits (33, 34). Hence, during *de novo* flagella synthesis, it begins at a minimum and increases generations later to strong propulsion by mature flagella (48). We hypothesized that the strong force produced by mature cells would separate sister cells and we investigated this using a motile strain (AW405) that is closely related to MG1655. In AW405, flagellation is repressed by glucose and induced in its absence. This is due to a mutation causing CRP-regulation of the flagella master regulator *flhDC* (49) and allows simple experimental control of flagellation. By fluorescently staining the flagella (50), we confirmed its presence required the absence of glucose (Fig. 3A & B). Next, tracking flagellated AW405 cells in glucose (−) conditions, we observed swimming cells and the separation of all sister cells after division (Fig. 3C; Movie S21), consistent with our hypothesis. Even after many generations, no folding, rosettes, or multicellular communities were observed. To verify the flagellum’s role, we tracked AW405 cells with a Δ*fliC* mutation in glucose (−) conditions and observed sister cells remained adhered at their poles, producing linear, single-cell-wide chains (Fig. 3D; Movie S22) identical to those of MG1655 Δ*fliC* and Δ*motA* cells. In agreement with the Δ*cheZ* data, these findings indicate that strong flagella propulsion separates sister cells and thereby abolishes rosette formation.

**Figure 3.**
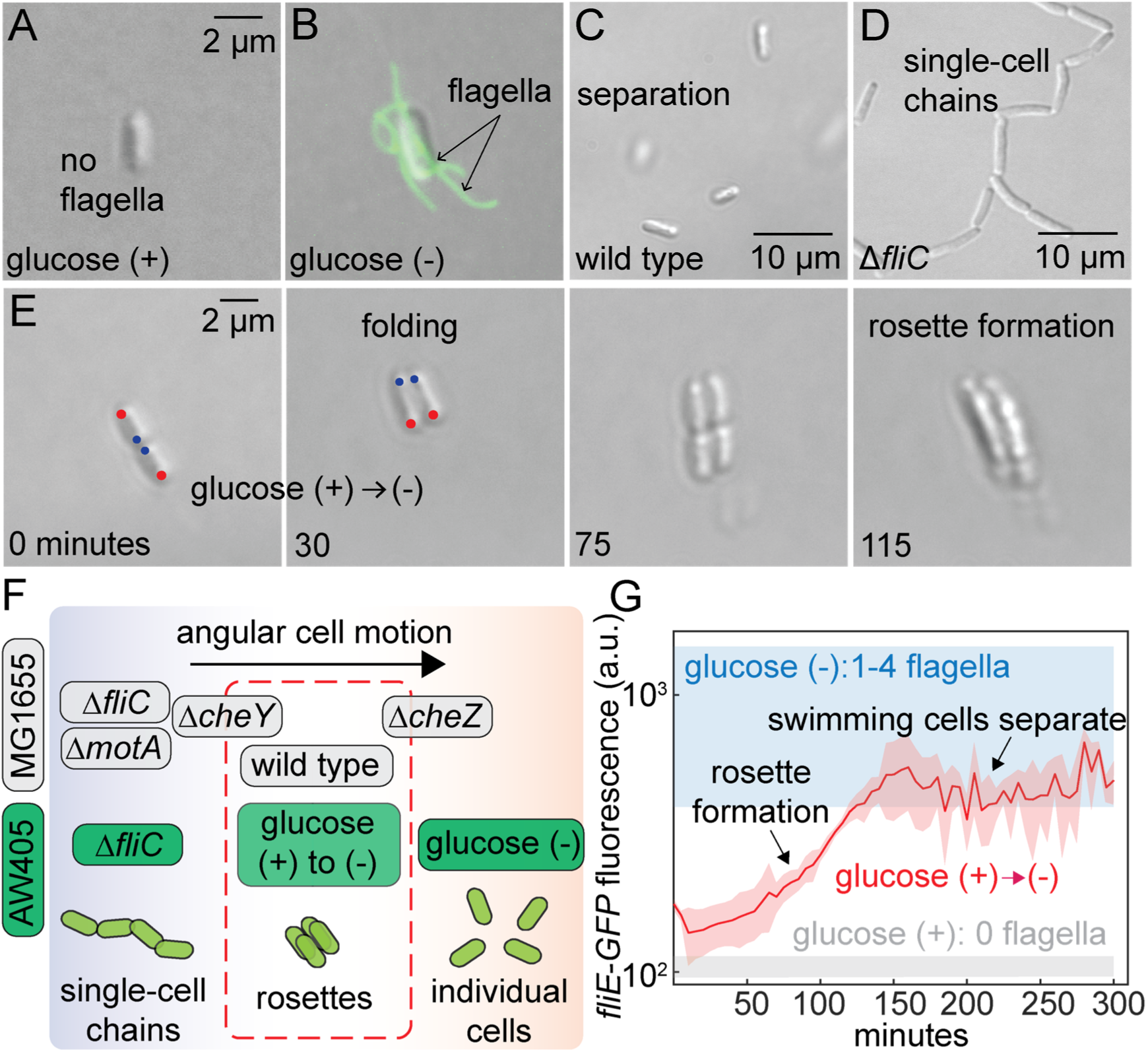
Rosette formation requires *de novo* flagella biosynthesis. (A) Representative micrographs of AW405 cells grown (A) with glucose and (B) without glucose and stained with Alexa Fluor 488 (green). Scale is indicated on micrographs. Micrographs from representative movies of (C) AW405 and (D) AW405 Δ*fliC* cells grown without glucose. (E) Micrographs from a representative movie of AW405 transitioned from glucose (+) to glucose (−) media. New (blue) and old (red) poles are indicated. (F) Diagram of the observed angular cell motion of strains and conditions used throughout this study and correspondence to community structure. Δ*cheY* and Δ*cheZ* cells produce heterogeneous outcomes and hence overlap depicted multicellular regions. (G) Cellular fluorescence (arbitrary units) from *fliE*-*GFP* promoter reporter during chain morphogenesis (red) in AW405 on transition from glucose (+) to (−) (5-minute resolution; mean ± standard deviation, n=3). Grey zone indicates *fliE*-*GFP* fluorescence in non-flagellated cells in glucose (+) media (see Fig. S37). Blue zone indicates *fliE*-*GFP* cellular fluorescence in flagellated cells (1-4 flagella per cell) in glucose (−) media (see Fig. S37). The approximate times of rosette formation and dissociation of single swimming cells are indicated by arrows.

Adding to the findings in defective-flagella mutants (Δ*fliC* nor Δ*motA*) (Fig. 2), the AW405 results suggest rosette formation requires intermediate forces, such as those produced during *de novo* flagella biosynthesis (48). Hence, we hypothesized that inducing *de novo* biosynthesis in previously non-flagellated AW405 cells might cause them to self-assemble into rosettes. To test this, we grew AW405 cells in glucose (+) media, transitioned them to glucose (−) to induce flagella biosynthesis, and tracked them by microscopy. In these experiments, sister cells folded at both the 2-cell and 4-cell stages, thereby producing quatrefoil rosettes identically to MG1655 (Fig. 3E; Movie S23). This finding indicates that inducing *de novo* flagella biosynthesis is sufficient to enable rosette formation in AW405 and supports the idea that sister cells must balance the opposing forces of adhesion and propulsion to properly reposition into rosettes (Fig. 3F). Balancing of *Ag43*-adhesion and flagella propulsion has been demonstrated in motility assays (51) and could be further characterized by similar studies at the cellular scale.

We next sought to investigate the dynamics of flagella biosynthesis during rosette formation. However, methods for flagella staining (50) have not been adapted for real-time tracking of *de novo* biosynthesis. Therefore, we characterized the transcriptional dynamics of the flagella pathway using a fluorescent promoter reporter for *fliE* (52), which is directly regulated by *flhDC* and the earliest gene induced during flagella biosynthesis (53). We validated this reporter in AW405 by comparing single-cell *fliE* expression to the number of flagella per cell, which was directly quantified by staining (50) and ranged from 1 to 4 (Fig. S37). We then tracked *fliE* induction dynamics in AW405 cells after transition from glucose (+) to glucose (−) in liquid culture, noting an exponential curve over several generations (Fig. S37). We next tracked *fliE* expression during rosette formation in devices upon transitioning from glucose (+) to glucose (−) as above (Fig. 3E). These experiments demonstrated that *fliE* expression was activated upon switching to glucose (−) and increased exponentially during rosette formation (Fig. 3G). At the time of rosette formation itself, *fliE* expression was between the fluorescence levels corresponding to 0 and 1 flagella per cell, suggesting an intermediate degree of flagella biosynthesis. At approximately the 16-cell stage, *fliE* expression plateaued at a fluorescence level corresponding to 1 flagella per cell (Fig. S37). In AW405, individual swimming cells separated from multicellular chains at later time points (∼32 cells). This behavior had not been observed in wild-type MG1655, and may result from the more robust activation of flagella biosynthesis in the AW405 strain. Together, these findings demonstrate that transcription of the flagella biosynthesis pathway is induced during rosette formation. Furthermore, the intermediate *fliE* activation suggests folding of rosette corresponds to intermediate flagella biosynthesis, and therefore force. In the future, the development of new real-time staining methods could enable more direct investigation of the flagellum’s role in rosette formation.

Though we had previously identified rosettes in *E. coli’*s multicellular life cycle (6), it was unclear if they were developmentally required for the subsequent morphogenetic stages. To test this idea we tracked wild-type, Δ*fliC*, and Δ*motA* communities long-term. We found that, even after many generations, both Δ*fliC* and Δ*motA* communities remained linear with cells adhered to one another at their poles (Fig. 4B, & Fig. S38-39) and did not display the well-regulated post-rosette morphogenesis of wild type (Fig. 4A) (6). In contrast to the stable organization of wild-type chains, Δ*fliC* and Δ*motA* communities were unstable and dissociated into smaller groups and individual cells, reminiscent of cells defective in type-1 fimbriae or curli (6). Moreover, Δ*fliC* and Δ*motA* communities poorly attached to cover glass (Fig. S38-39; Movie S24), reminiscent of PGA-defective cells (6). These results show that disabling rosette formation impairs subsequent morphogenetic stages. To gain additional insights into the developmental consequences of rosette formation, we tracked the expression of the alternate sigma factor σ^38^ (encoded by *rpoS*) during chain morphogenesis using a fluorescent promoter reporter (52). σ^38^ is a transcriptional master regulator and controls a variety of genes in biofilms, including those involved in stress tolerance and metabolism (54). We found that *rpoS* expression was strongly induced in wild-type chains between the ∼200-cell and ∼1000-cell stages prior to surface attachment (Fig. 4C). Despite identical growth conditions, rosette-disabled Δ*fliC* communities did induce *rpoS* expression (Fig. 4C). Together, these findings indicate that rosette formation is a developmental requirement for *E. coli*’s facultative multicellular life cycle, and may also be critical to biofilm formation.

**Figure 4.**
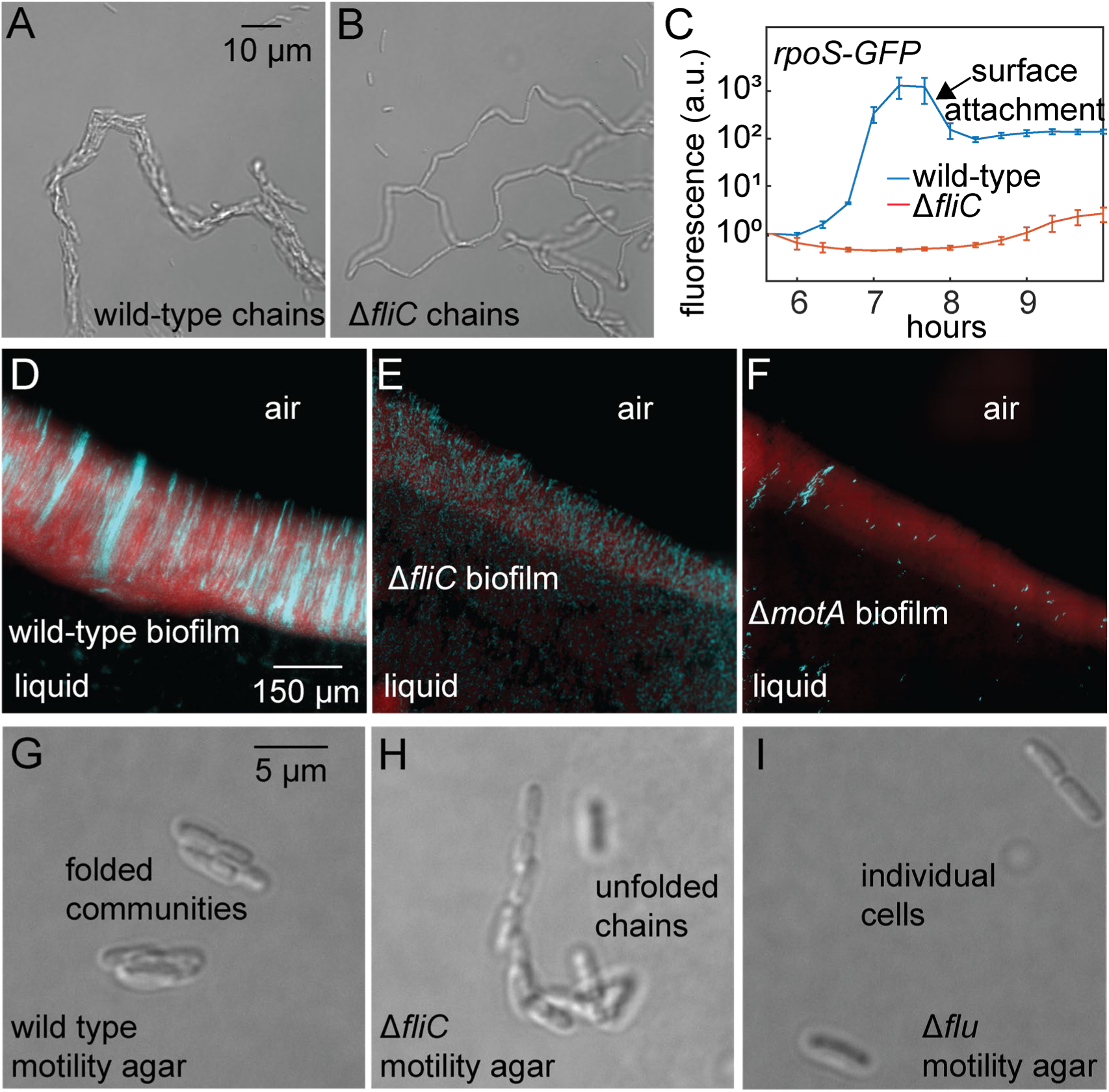
Developmental consequences and generality of rosette formation. Micrographs at 5 hours from representative movies of (A) wild-type and (B) Δ*fliC* community formation (scale is indicated). (C) Cellular fluorescence (arbitrary units) from *rpoS*-*GFP* promoter reporter during chain morphogenesis in wild type (blue) and Δ*fliC* (red) (20-minute resolution; mean ± standard deviation, n=3). The approximate time of chain attachment to surfaces is indicated. Micrographs of hydrostatic biofilms formed by (D) wild-type, (E) Δ*fliC*, and (F) Δ*motA* cells. Cells contained either mScarlet (red) or GFP (cyan) at a 10:1 ratio (scale, as well as air and liquid sides of biofilms, are indicated). Example micrographs of (G) wild-type, (H) Δ*fliC*, and (I) Δ*flu* cells after ∼2 hours on motility agar (2-layer 0.25% agar; scale is indicated; see also Fig. S40).

We have previously shown that *E. coli*’s rosette-initiated chain morphogenesis generates hydrostatic biofilms (6). Hence, we hypothesized that Δ*fliC* and Δ*motA* mutants, which disable rosette formation, would reduce surface attachment and disrupt the clonal-chain architecture of *E. coli* biofilms (6). To test this, we grew hydrostatic wild-type, Δ*fliC*, and Δ*motA* biofilms and imaged them at the cellular scale (7). Consistent with our hypothesis, these mutations reduced biofilm volume and abolished the parallel-aligned, clonal-chain structure of wild-type biofilms (Fig. 4D-F). Moreover, Δ*fliC* and Δ*motA* biofilms contained short unfolded chains of cells adhered at their, structurally consistent with the lack of flagella propulsion and rosette formation. These findings further demonstrate the developmental consequences of rosette formation and provide a new rationale for the requirement of flagella to hydrostatic *E. coli* biofilms (37, 38).

We next considered that flagella-assisted rosette formation might be general in hydrostatic environments. Aside from biofilm assays, hydrostatic conditions are commonly studied at bulk-scale in motility assays (55, 56). The “swimming agar” used in such assays (∼0.2-0.3% agar) provides cells sufficient freedom of motion to swim across the surface. On the other hand, “colony agar” (e.g., >1.5% agar for many species) limits free motion and forces growth within localized colonies. We reasoned that, if rosette formation was general to hydrostatic environments, then we should observe them during *E. coli* growth on swimming agar. Conducting standard motility assays (51, 57) and imaging after ∼2 hours, we observed freely-moving rosette-like communities and multicellular chains in the regions where cells had been initially deposited, suggesting rosette formation can occur on swimming agar (Fig. 4G; Movies S25-27). In these experiments, we also observed examples of cellular folding in real time (Movies S28-29). To investigate the mechanism of community folding on swimming agar, we performed similar experiments with Δ*fliC* and Δ*flu* (*Ag43*) strains. We observed that Δ*fliC* cells were mostly adhered at the poles in linear, unfolded chains, demonstrating the flagellum is required for folding on swimming agar (Fig. 4H; Movie S30). Moreover, Δ*flu* cells were predominately individuals and were rarely observed in aggregates (Fig. 4I; Movie S31). These findings are in agreement with a previous study demonstrating balancing between *Ag43* adhesion and flagella propulsion on motility plates (51). Moreover, they indicate the formation of rosettes on swimming agar and the requirement for *Ag43* adhesion and flagella propulsion. Additionally, the imaging technique we have developed could be optimized and extended to further investigate the genetics and dynamics of bacterial multicellular behavior in hydrostatic environments.

## Discussion

This study has revealed *E. coli* forms rosettes by a simple mechanism, flagella-assisted repositioning of sister cells. Flagella propulsion produces an angular random walk between sister cells (Fig. 1-3), thereby folding them in parallel alignment at the 2- and 4-cell stages (Fig. 1). The synchronization of cell division events further explains the robustness of quatrefoil rosettes and why alternatives like 3-, 5-, or 6-cell rosettes were not observed. We demonstrated that rosettes are required for subsequent morphogenesis of multicellular chains, *rpoS* expression, and formation of clonal-chain biofilms (Fig. 4). Consistent findings were made in diverse experimental conditions, including devices (Fig. 1-3), biofilms (Fig. 4), and swimming agar (Fig. 4), suggesting that rosette formation may be a general behavior in hydrostatic environments and has a straight-forward dependence on mechanical constraints. The small forces produced by mature flagella are sufficient to separate sister cells (Fig. 3), indicating bulk-scale forces, like mechanical shaking for culture agitation, would necessarily block rosette formation and would thereby effectively explain why fluid flow impairs commensal *E. coli* biofilm formation (58). On the other hand, forcing cells against surfaces, such as colony growth on agar or on cover glass in flow cells, constrains cell motion and blocks repositioning into rosettes. Mechanically speaking, hydrostatic conditions would then be necessary for commensal *E. coli* rosette formation and subsequent biofilm morphogenesis.

This study also updates *E. coli*’s facultative multicellular life cycle (6), which has now provided roles for many of the genes required for hydrostatic biofilm formation (38). Adding to those outlined in the introduction, our current findings revealed new multicellular behaviors: induction of flagella biosynthesis at the 2-cell stage leads to cellular repositioning at the 2- and 4-cell stages to produce rosettes; and expression of *rpoS* is induced at ∼200-cell stage. The active mechanism of rosette formation and its requirement for subsequent multicellular behaviors further support the idea that *E. coli*’s facultative life cycle is an organized developmental process with regulated stages (illustrated in Fig. 5). In the future, the generative structural interactions could be further validated by new real-time staining techniques. This pathway may be specific to *E. coli,* perhaps reflecting its natural habitats, but other bacteria perform similar phenomena. *Bacillus subtilis*, for example, can produce biofilms by multicellular chain morphogenesis (59) which is maintained by ECM production (60) and has genetically-regulated stages (61), though rosettes have not been reported. As reported here, *E. coli*’s mechanism of rosette formation is currently unique as the rosettes described in other species either propagate through binary fission (31) or result from the entanglement of flagella without enclosing an internal space (28–30).

**Figure 5.**
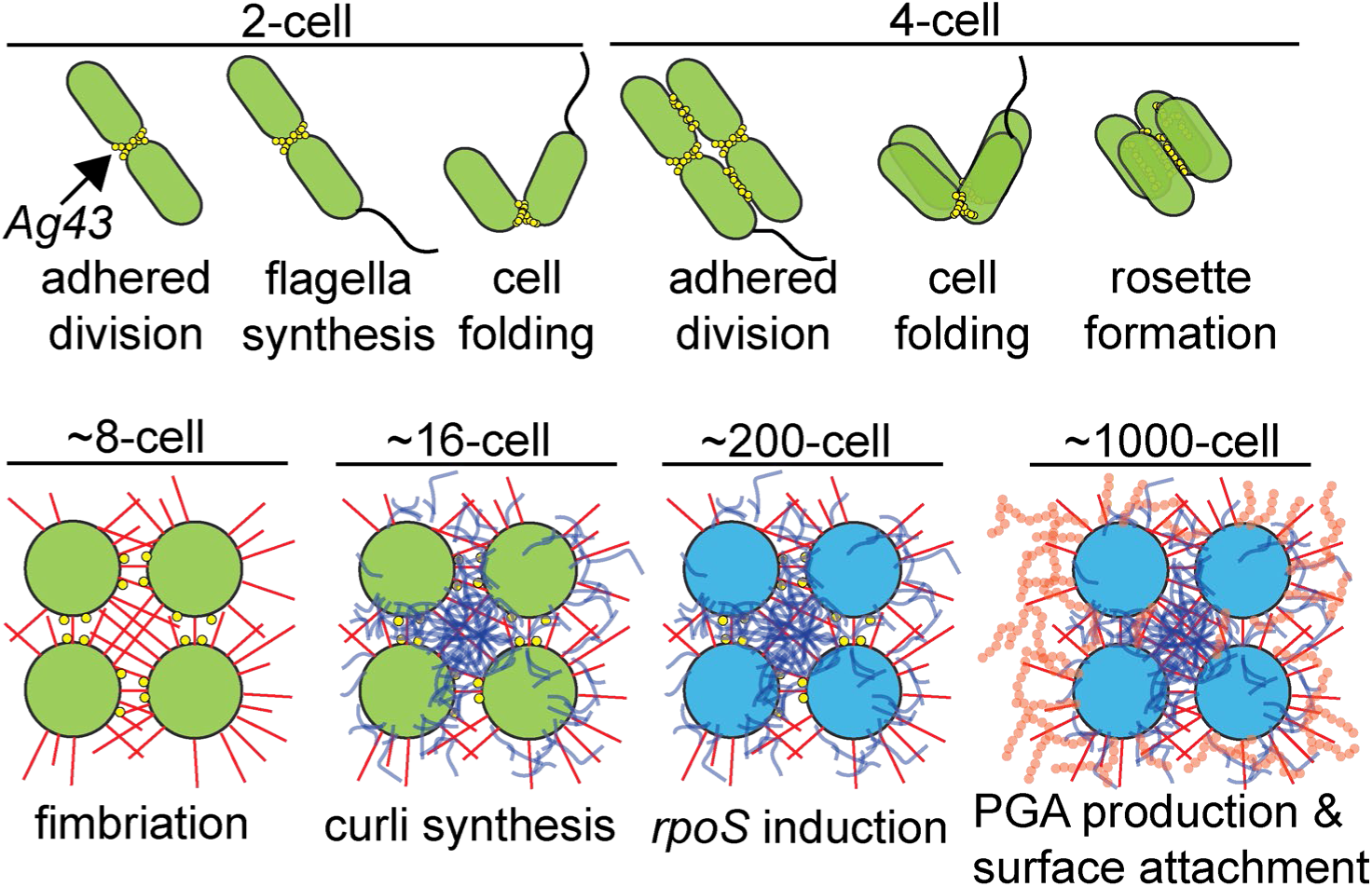
Stages of *E. coli* multicellular morphogenesis in hydrostatic environments. The data presented here and in our previous study (6), together with past characterization of ECM-comprising adhesins, support the following working model. 2-cell: self-recognizing *Ag43* enables adhesion of sister cells after the first division event. Cells induce *de novo* flagella biosynthesis, and the propulsion of the growing flagella tail causes sister cells to perform cell folding by an angular random walk to align side by side. 4-cell: sister cells adhere after the second division event and again perform cell folding to align in parallel and produce rosettes with quatrefoil configuration. Rosettes transition into chains, whose square cross-sectional arrangement is approximately maintained. ∼8-cell stage: radial-projecting extracellular polymer type-1 fimbriae (indicated in red) controls chain stability. ∼16-cell stage: extracellularly polymerizing curli (indicated in blue) maintains cellular positioning. ∼200-cell stage: transcriptional master regulator σ^38^ (encoded by *rpoS*) is induced (indicated by cells turning blue). ∼1000-cell stage: extracellular polysaccharide poly-b-1,6-N-acetyl-D-glucosamine (PGA, indicated in orange) is produced and attaches multicellular communities to surfaces. The representations of adhesins are not to scale and have yet to be confirmed by electron microscopy.

Moreover, this study is the first report of clonal rosette formation by cellular repositioning in bacteria. In principle, the same behavior underlies simple eukaryotic multicellularity and is hypothesized as a key step in the evolution of complex organisms (4). Similar to *E. coli*, *S. rosetta* cells with mature flagella separate and those lacking flagella adhere in unfolded linear chains (23). Hence, our study demonstrates that clonal self-organization of developmental rosettes is a cross-kingdom behavior and may represent a common design principle of multicellularity, irrespective of an organism’s genomic complexity or evolutionary history. In turn, this principle suggests rosette self-assembly might provide a foundational stage for programming synthetic multicellular behaviors, a goal for *E. coli* (62) since early on in synthetic biology (63, 64). Though *E. coli* lacks complexity relative to even the simplest eukaryotes, rosette formation may provide a basis for engineering or evolving increasingly complex behavior in a tractable unicellular organism.

This study additionally suggests rosettes might be targeted to control *E. coli*’s natural multicellular behaviors, including the formation of biofilms. Hydrostatic biofilms depend on rosette formation (Fig. 4) which in turn relies on the balance of sister-cell adhesion and flagella propulsion (Fig. 3). Tuning this balance could potentially be used to modulate the biofilm capacity of commensal *E. coli* strains, which form multicellular communities (65) in the ∼1 mm-thick intestinal mucosa (66) as members of the microbiome (67). Alternatively, flagella motion could be blocked to therapeutically inhibit *E. coli* biofilms, which contribute to urinary-tract infections (68) and are highly tolerant to antibiotics (69). This tolerance depends on expression of biofilm-related genes, including those regulated by *rpoS* (70). Hence, as rosettes are critical to adaptive gene expression (Fig. 4), disrupting their formation might also enhance antibiotic susceptibility. Such rosette-targeting approaches could be combined with newly-discovered antibiotics (71–73) or complement mechanism-based improvement of existing antibiotics (74, 75), such as the metabolic potentiation of aminoglycosides (76–79).

## Materials and Methods

### Strains, Plasmids, and Media

*Escherichia coli* MG1655 was the primary strain used in experiments. *E. coli* AW405 strains were also used where specified. To facilitate computational image processing and analysis and also biofilm imaging, *E. coli* cells were transformed with pUA66-p*ompC*::*gfp* (green) (52) or pEB2-mScarlet (red) (80). These plasmids constitutively express fluorescent proteins and had no observable effects on cellular behaviors. Additionally, pUA66-p*fliE*::*gfp* and pUA66-p*rpoS*::*gfp* (52) were also transformed into MG1655 and studied where described. The Δ*fliC*, Δ*motA*, Δ*cheZ*, Δ*cheY*, and Δ*flu* strains were created by transducing Δ*fliC*::kan(kanR), Δ*motA*::kan(kanR), Δ*cheZ*::kan(kanR), Δ*cheY*::kan(kanR), and Δ*flu*::kan(kanR) respectively from the KEIO Collection (81) using *P1vir* in *E. coli* MG1655 following standard methods (82), and were cured with pCP20 (83). All strains were cultured in LB, Miller broth (Difco) at 37°C and with shaking at 300 RPM, in 14 mL Falcon Round-Bottom Test Tubes with aerating caps (VWR), and in light-insulated shakers. LB lacks significant glucose and was therefore used as a glucose (−) condition. Glucose (+) experiments used Neidhardt Supplemented MOPS Defined Medium (EZ-rich; Teknova) (84). When necessary, plasmids were maintained by supplementing media with 50 μg/mL of kanamycin.

### Cell Tracking in Hydrostatic Devices

LB agarose pad devices were built using 2% agarose, following previously described methods (6). 3 µL exponential phase cells were added to agarose pads, which were immediately covered with cover glass without allowing them to dry. This generated a liquid space ∼0.75 cm x 0.75 cm that we approximated as ∼5-10 µm deep based on the distance between the focal planes of the cover glass and agarose. These devices were developed to allow observation of the free motion of cells in a hydrostatic environment. For glucose transition experiments, *E. coli* AW405 cells were first cultured overnight in Neidhardt Supplemented MOPS Defined Medium (EZ-rich; Teknova) and then loaded onto LB-2% agarose pads for cell tracking. Dynamic bright-field imaging was typically conducted at 2-second resolution. Dynamical fluorescent imaging for motion tracking was performed at 30-second resolution (instead of 2-second) to avoid the confounding effects of exposing bacteria to high-intensity blue light. Imaging for fluorescent reporters (pUA66-p*fliE*::*gfp* and pUA66-p*rpoS*::*gfp*) was less frequent to reduce photo-bleaching and is specified in legends.

### Biofilm Imaging

Biofilm experiments were performed as previously described (6, 7). Wells in 96-well plates were loaded with 200 mL of fresh LB with overnight cultured cells (∼10^8^ CFU/mL). Cultures were a 1:10 mixture of cells containing pUA66-p*ompC*::*gfp* (green) and pEB2-mScarlet (red). Cover glass pieces were placed in each well, perpendicular to the air-liquid interface in order to allow for biofilm formation. Plates were then incubated without shaking at 37°C for 16 hours to allow biofilm formation to occur. After incubation, cover glass pieces were removed from the wells and placed between a glass slide and a fresh piece of cover glass in order to image by microscopy. This method allows high-magnification imaging of hydrostatic biofilms grown following a standard protocol (10).

### Motility Assay Imaging

Swimming agar is fragile and difficult to remove from a petri dish without damage. To enable handling of swimming agar, we created two layer plates by adapting standard procedures (57). The bottom layer was made from 2% agar and provided a stable base for a top layer of 0.25% swimming agar. This ensured cells spotted on these plates would encounter an identical hydrostatic surface as in classical motility assays and would also have the same environmental nutrients concentrations. We confirmed comparable bulk-scale motility on 2-layer (2% and 0.25% agar) plates and classical swimming agar plates (0.25 % agar) by incubation for 18 hours at 37°C (Fig. S40) following previous methods (51). For imaging directly on motility plates, we first poured a 12 mL layer of molten 2% LB agar in petri dishes and allowed them to cool overnight. The day of motility experiments, we poured 8 mL of molten 0.25% LB agar (at 50°C) on top of the 2% layer and cooled. 2-layer plates were pre-warmed at 37°C, then 10 μL of exponentially growing cell suspensions diluted to ∼10^5^-10^6^ CFU/mL were spotted in the middle. Plates were then incubated face up for 1-2 hours at 37°C. After incubation, a ∼1 cm x 1 cm square containing the location of the initial 10 μL spot was cut in the agar using a scalpel. The remaining agar was removed from around this square. Cover glass was placed on top of this square, resting on the hydrostatic surface of the swimming agar. Immersion oil was added to the cover glass, then the agar and cover glass were carefully inverted and placed directly in a microscope sample holder and immediately imaged. As cells and communities were in motion, capturing them required recording videos in bright-field DIC.

### Microscopy

Images were acquired using a Leica DMi8 microscope equipped with a DIC HCPL APO 63X oil immersion objective (with 1.6x magnification changer), Hamamatsu ORCA-Flash 4.0 camera, and Lumencor Spectra-X light engine. Devices for imaging were secured in a microscope-stage-top incubator (Tokai-Hit, STX Stage Top Incubator Temp and Flow; STXF-WSKMX-SET), and the incubator and the microscope objective were maintained at 37°C prior to and throughout all experiments. Time courses were collected in an automated manner using the Leica LasX software at specified time intervals. Imaging was performed using differential interference contrast (DIC) and fluorescence channels. Excitation and emission for fluorescence microscopy was performed at 470 nm and 500-550 nm for green fluorescence and 510 nm and 592-668 nm for red fluorescence respectively. AW405 flagella were stained as previously described (50) with Alexa Fluor 488. Micrograph examples and movies were generated using Leica X software.

### Image Analysis

Fluorescent images of cells were analyzed using built-in Matlab functions. Briefly, raw images were cropped to include only one sister-cell pair or multicellular community then were segmented using the *imbinarize* function. Centerlines of cells were then determined using the *bwskel* function and used to calculate the angle θ between sister cells. *imbinarize* was also used for the transcriptional reporter data (*fliE* and *rpoS*), and the average pixel intensity of the segmented area was calculated to determine cellular fluorescence. Bright-field DIC images of cells were first processed in ImageJ to manually mark the poles for both the sister cells. Coordinates for these pole positions were then loaded into MATLAB for calculating motion. θ was determined as the 2D angle created by the pole positions of two sister cells, and these values were also used to determine the angular velocity. Ф represented the angle between a cell and the imaging surface, in the z direction (toward the camera). This was imputed by trigonometry from the calculated length of the 2-D projection of the cell (the xy distance between a cell’s two poles when it was at an angle to the xy plane) and its calculated maximum length immediately prior to folding. As folding occurred within seconds, negligible cell growth (which has a doubling time ∼30 minutes) occurs, and cell length was treated as constant when calculating Ф. Strokes were identified as motion between consecutive 2-second images that exceeded 0.8 μm. δ was defined as the angle in 3D space between the vectors created by two consecutive strokes. Figures were generated in Matlab and arranged and formatted using Adobe Illustrator.

### Replicates and Statistical Analysis

Each replicate corresponds to an independent sister-cell pair or chain community, tracked over time. Replicates were chosen as representative examples from 3 or more separate experiments, and their number (n) is listed in figure legends. Mean and standard deviations are calculated where indicated. Micrographs and images are representative examples from replicated experiments.

## Acknowledgements

We are grateful to Howard C. Berg and Karen Fahrner for sharing AW405 strains and Philippe Cluzel for sharing pEB2-mScarlet. This study was supported by an NIH Director’s Early Independence Award (DP5OD019792) to Kyle R. Allison.

