## Supporting Information for "*Escherichia coli* self-organizes developmental rosettes"

**This PDF file includes:**

Supplementary Figures S1-S40

Legends for Movies S1-S31

**Other Supporting Materials for this manuscript:**

Movies S1-S31

### Supporting Figures

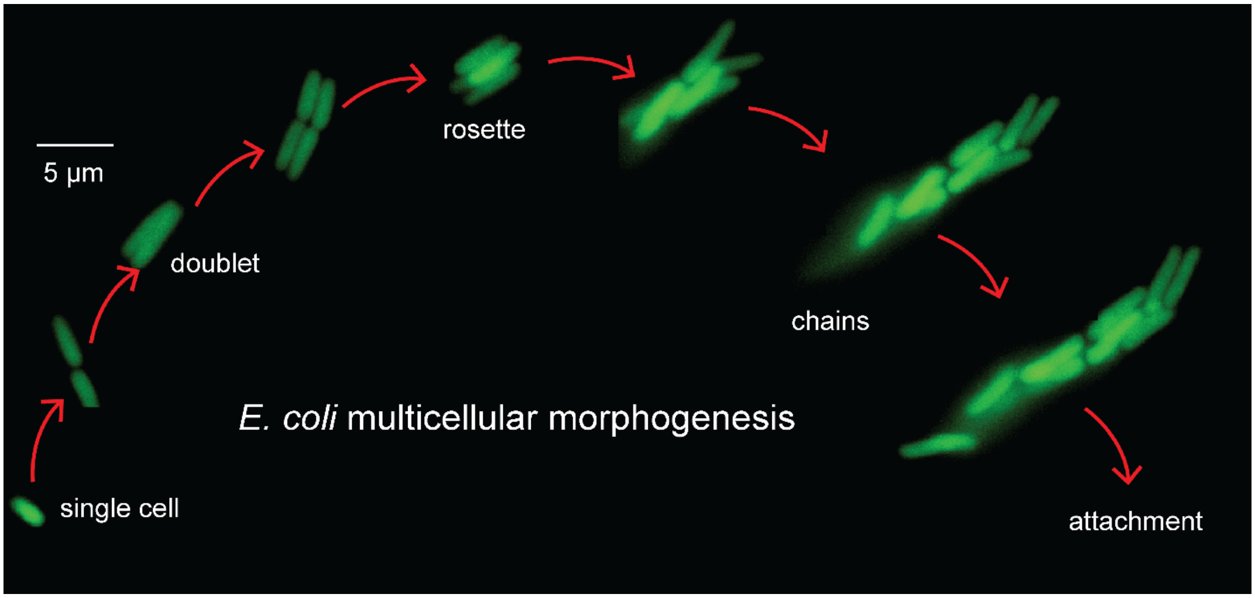


**Figure S1**. Example dynamic trajectory of rosette-initiated multicellular morphogenesis in *E. coli* containing green fluorescent protein (GFP). Scale indicated on micrographs.

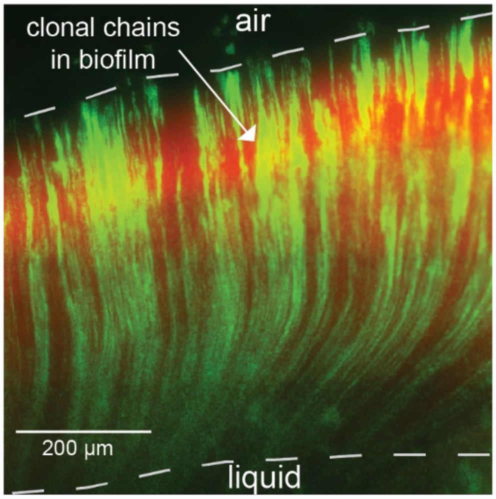


**Figure S2.** Direct visualization of *E. coli* biofilm imaged in multi-fluorescence. Biofilm was grown for 24 hours from a 1:10 mixture of cells containing GFP (green) and mScarlet (red) on cover glass, depicted in Figure 2C. Scale indicated on micrograph.

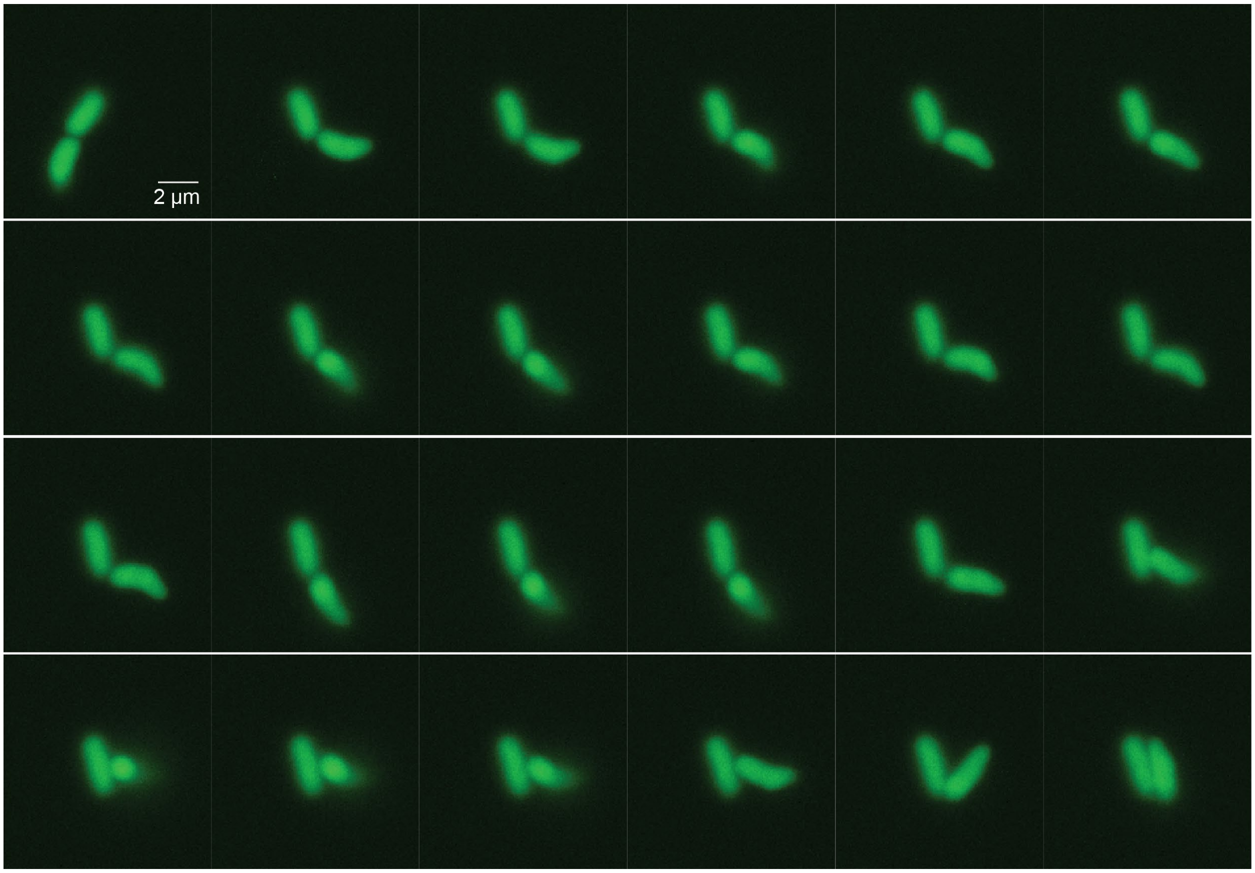


**Figure S3.** Example of cell folding in *E. coli* wild-type cells containing green fluorescent protein (GFP) (pUA66-p*ompC*::*gfp*) at 37°C. Exponential phase cells, grown in LB media, were loaded onto microfluidic device and imaged every 30 seconds. Each micrograph is 30 seconds apart and scale indicated on micrographs.

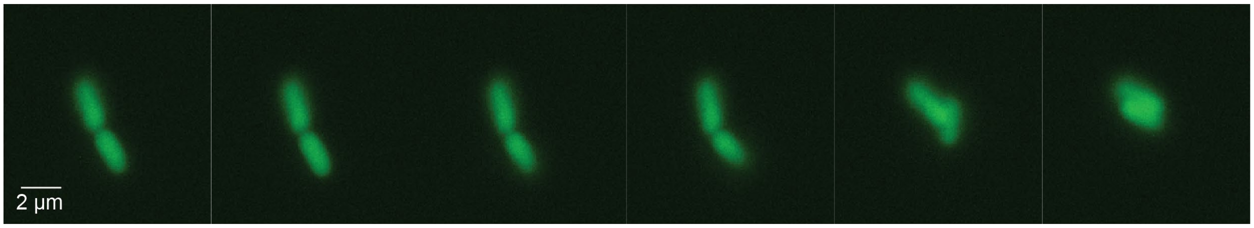


**Figure S4.** Example of cell folding in *E. coli* wild-type cells containing green fluorescent protein at 37°C. Each micrograph is 30 seconds apart and scale indicated on micrographs.

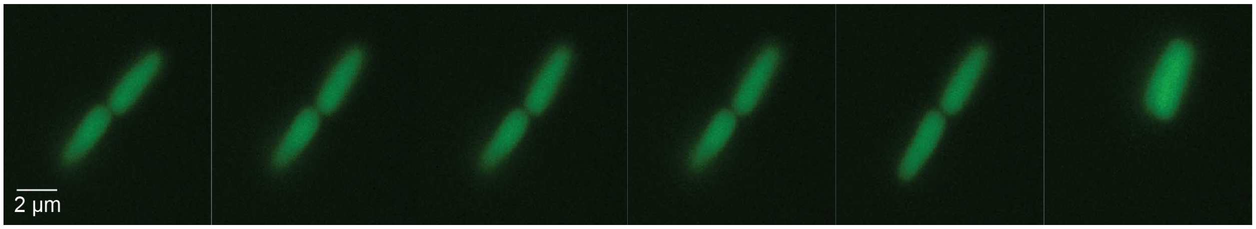


**Figure S5.** Example of cell folding in *E. coli* wild-type cells containing green fluorescent protein at 37°C. Each micrograph is 30 seconds apart and scale indicated on micrographs.

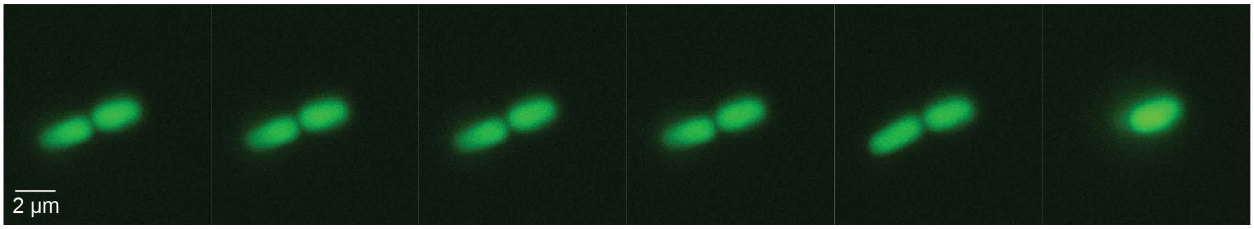


**Figure S6.** Example of cell folding in *E. coli* wild-type cells containing green fluorescent protein at 37°C. Each micrograph is 30 seconds apart and scale indicated on micrographs.

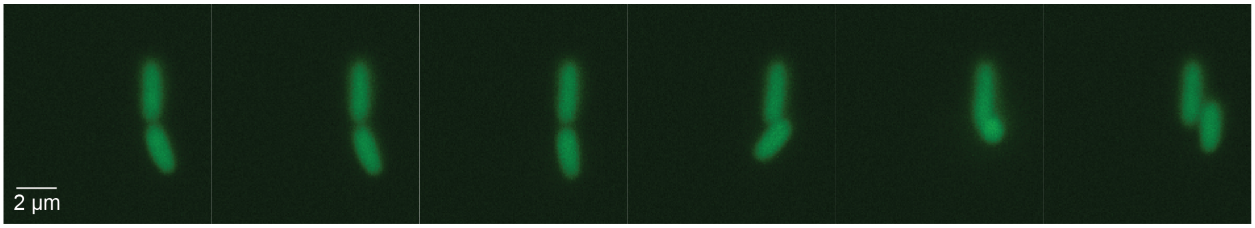


**Figure S7.** Example of cell folding in *E. coli* wild-type cells containing green fluorescent protein at 37°C. Each micrograph is 30 seconds apart and scale indicated on micrographs.

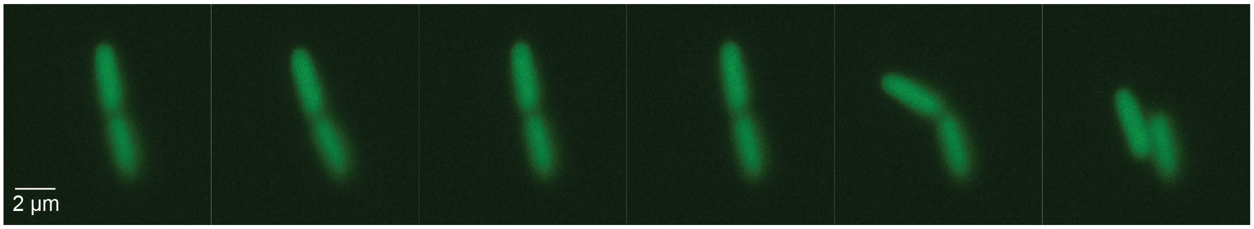


**Figure S8.** Example of cell folding in *E. coli* wild-type cells containing green fluorescent protein at 37°C. Each micrograph is 30 seconds apart and scale indicated on micrographs.

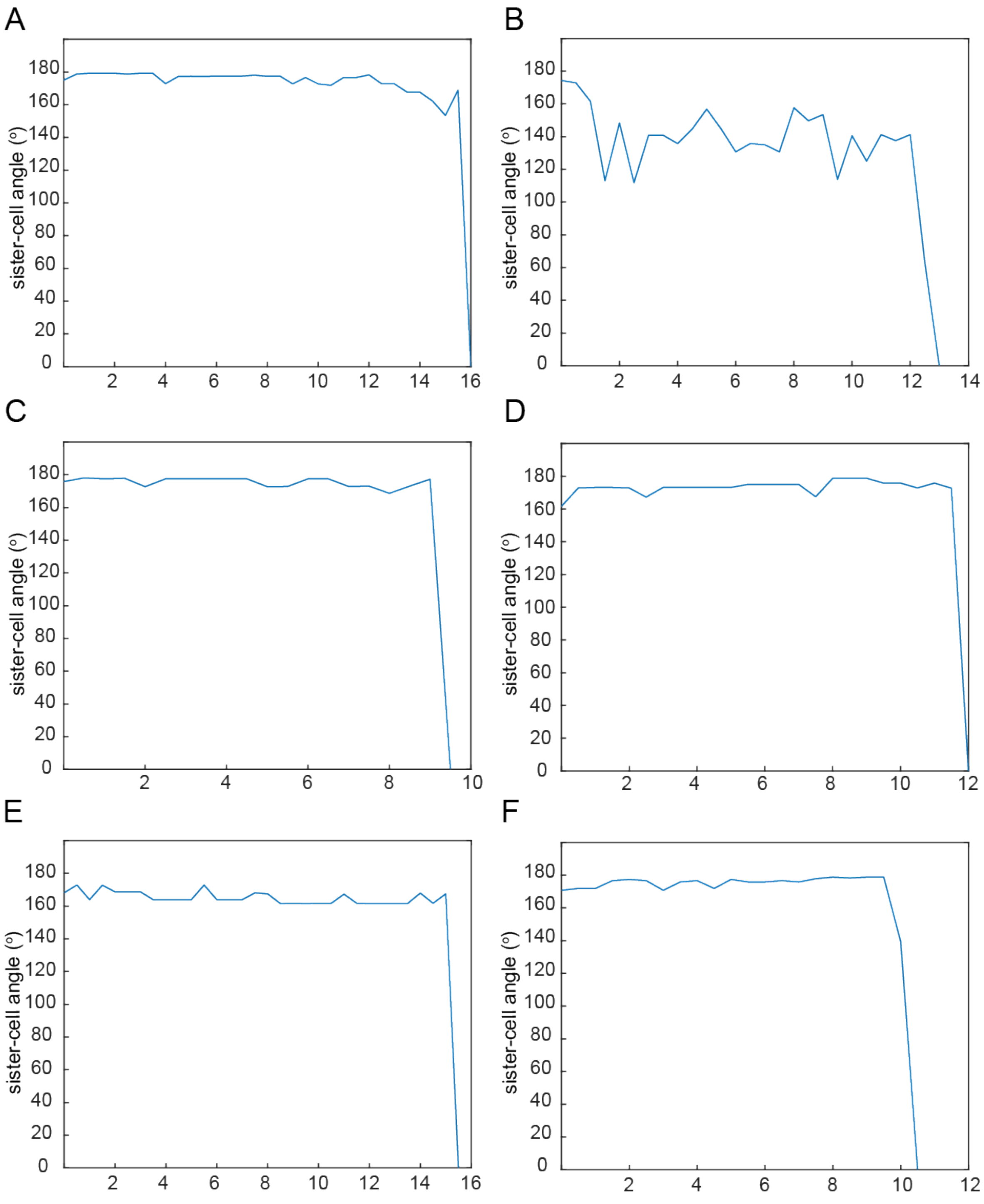


**Figure S9.** Individual examples of sister-cell angle dynamics during cellular folding (30-second resolution).

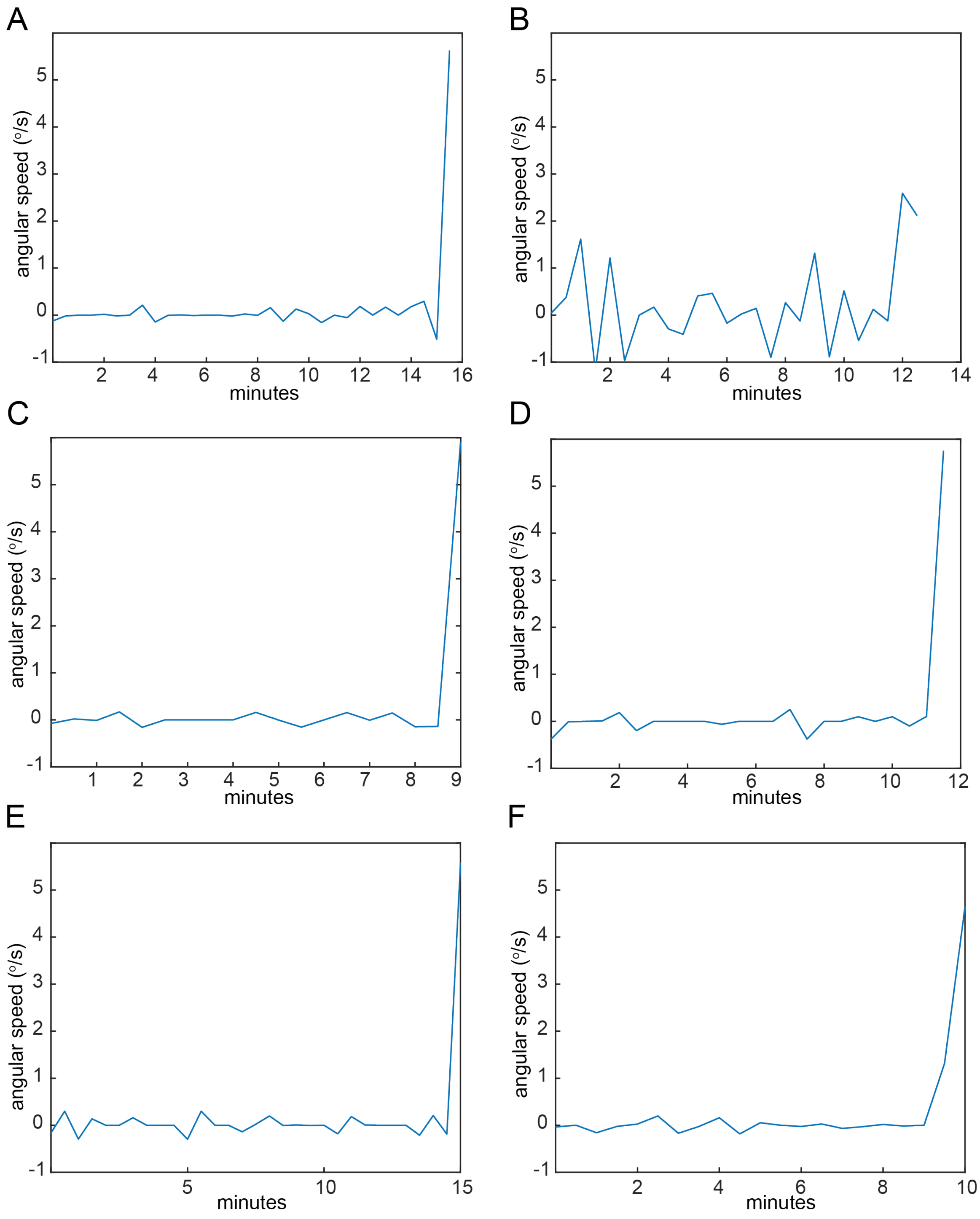


**Figure S10.** Individual examples of sister-cell angular speed during cellular folding (30-second resolution).

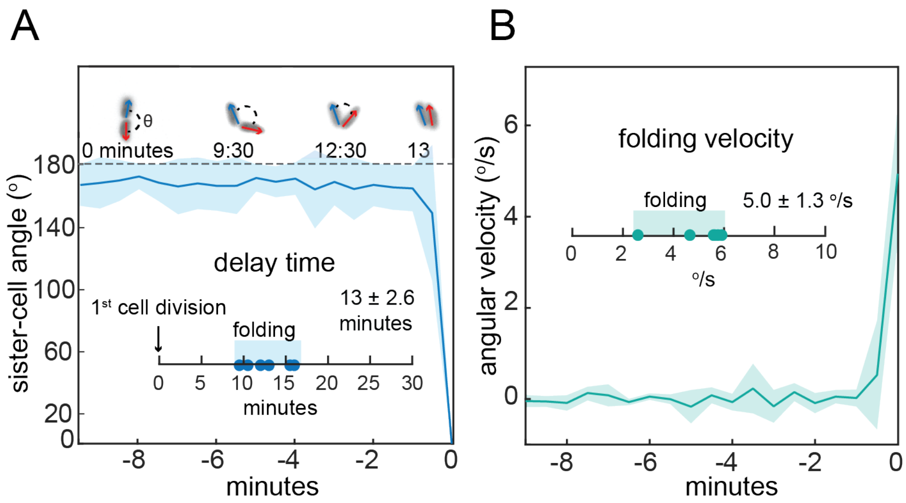


**Figure S11.** (A) Dynamics of sister-cell angle at 2-cell stage during cellular folding (30-second resolution; mean ± standard deviation, n=6). Inset: delay times between cell division and the completion of cell folding. (B) Dynamics of sister-cell angular speed at 2-cell stage during cellular folding (30-second resolution; mean ± standard deviation, n=6). Inset: Maximum folding speed per cell trajectory.

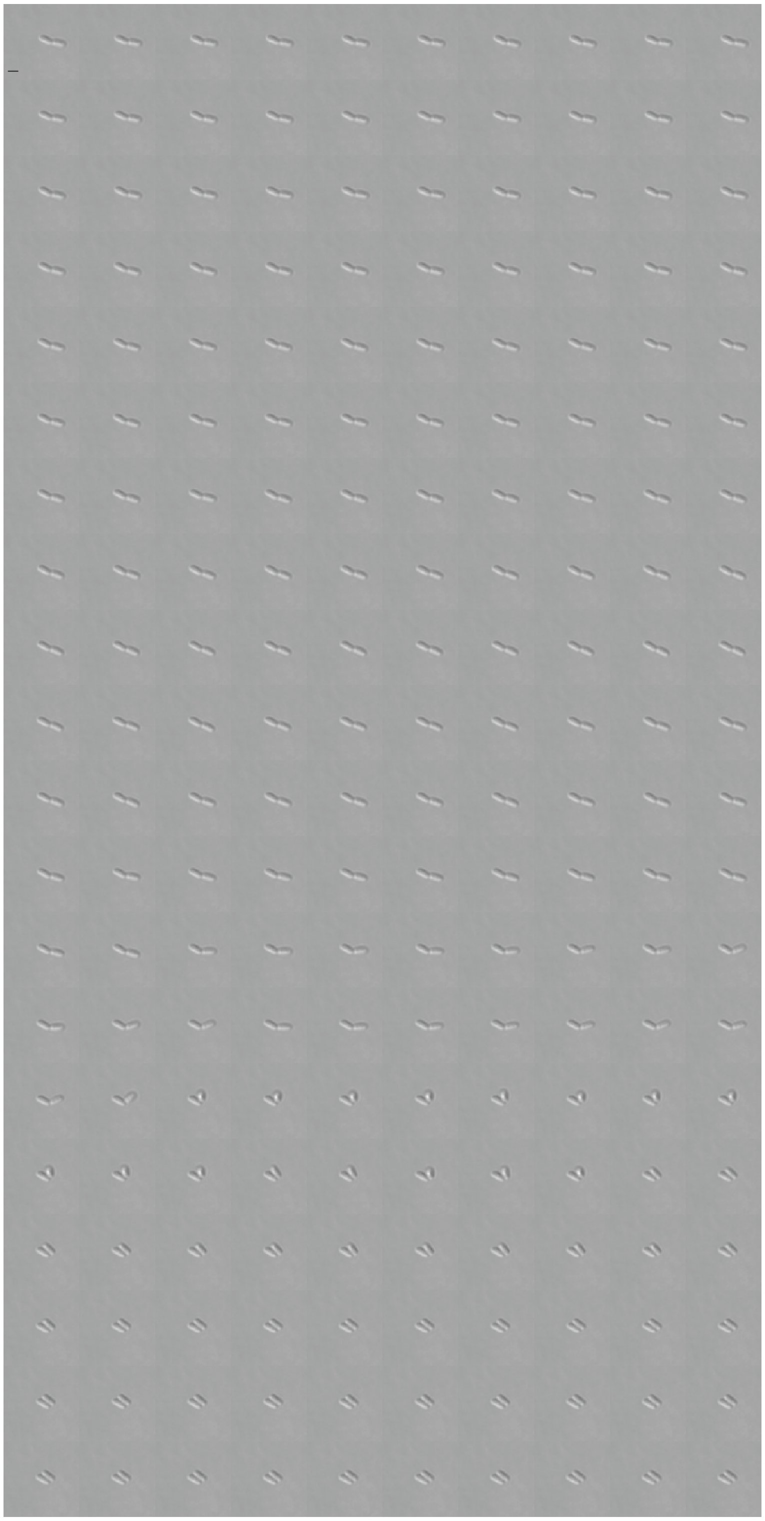

**Figure S12.** Example of cell folding in *E. coli* wild-type cells at in DIC at 37°C. Each micrograph is 2 seconds apart and scale bar on first micrograph represents 2 μm.

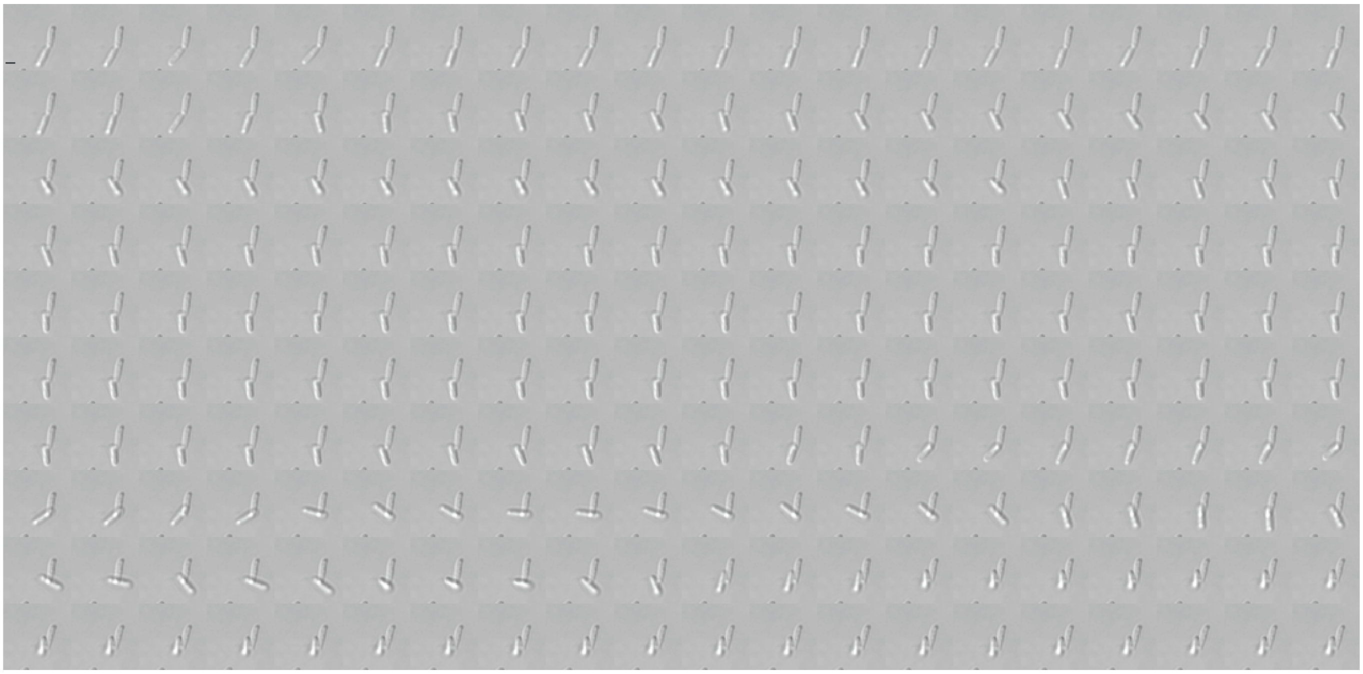

**Figure S13.** Example of cell folding in *E. coli* wild-type cells at in DIC at 37°C. Each micrograph is 2 seconds apart and scale bar on first micrograph represents 2 μm.

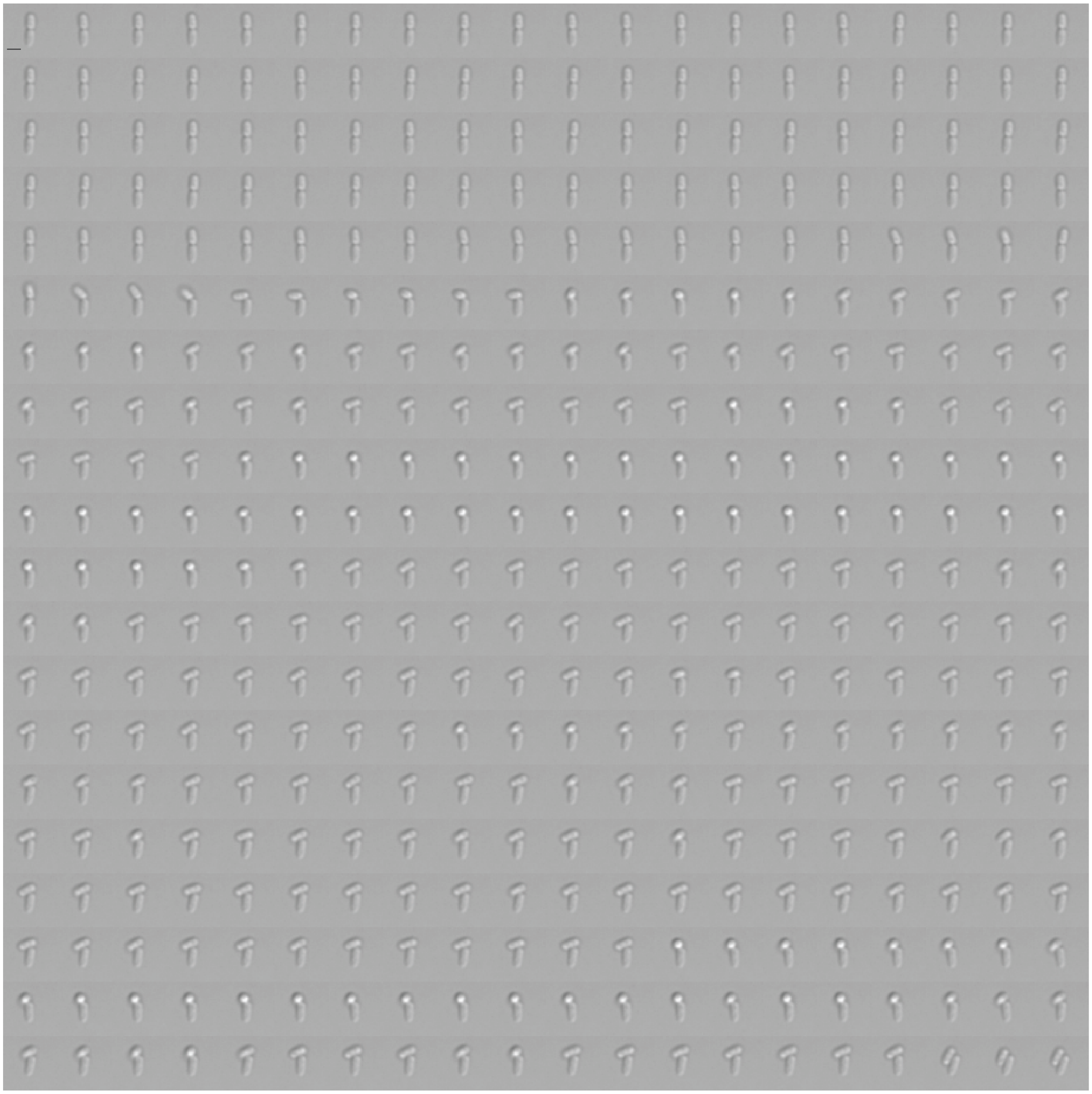

**Figure S14.** Example of cell folding in *E. coli* wild-type cells at in DIC at 37°C. Each micrograph is 2 seconds apart and scale bar on first micrograph represents 2 μm.

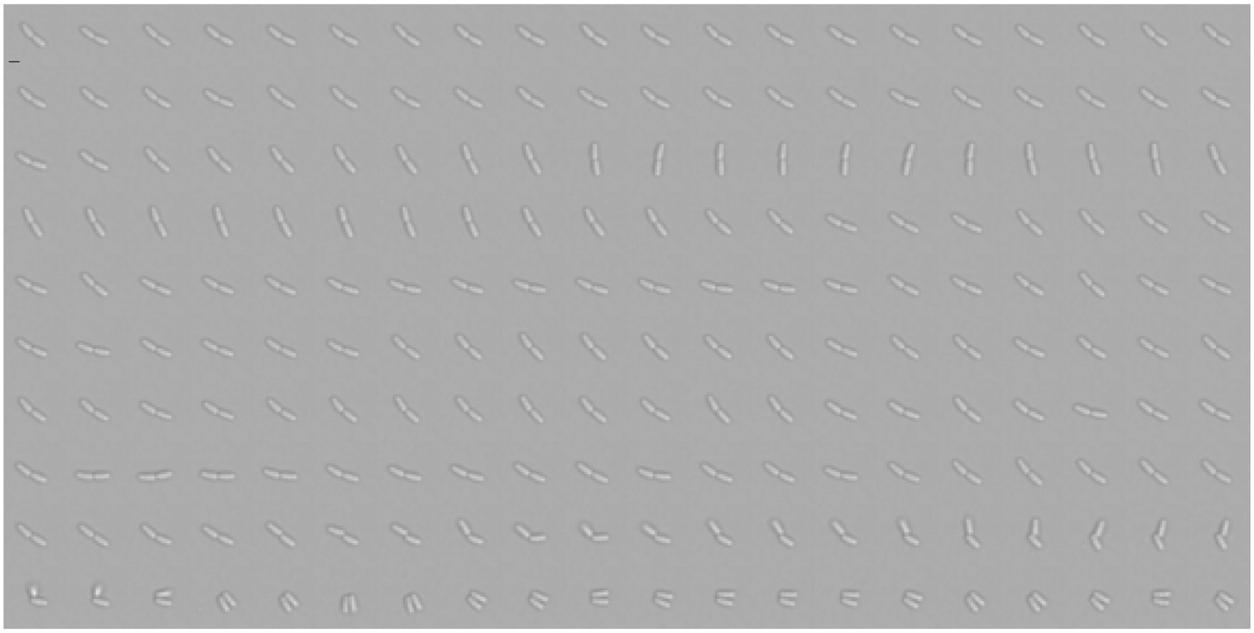

**Figure S15.** Example of cell folding in *E. coli* wild-type cells at in DIC at 37°C. Each micrograph is 2 seconds apart and scale bar on first micrograph represents 2 µm.

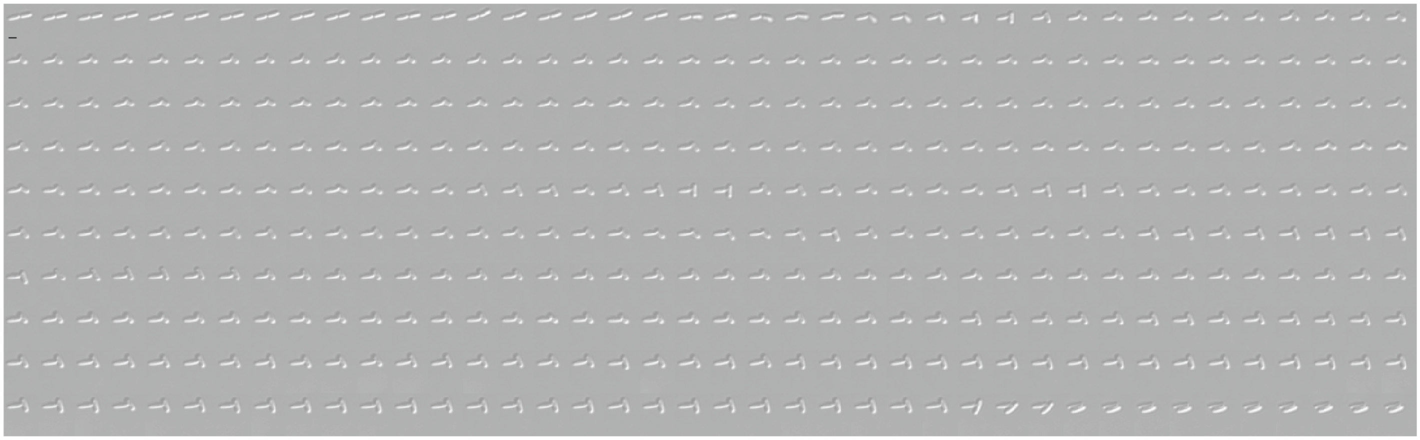

**Figure S16.** Example of cell folding in *E. coli* wild-type cells at in DIC at 37°C. Each micrograph is 2 seconds apart and scale bar on first micrograph represents 2 μm.

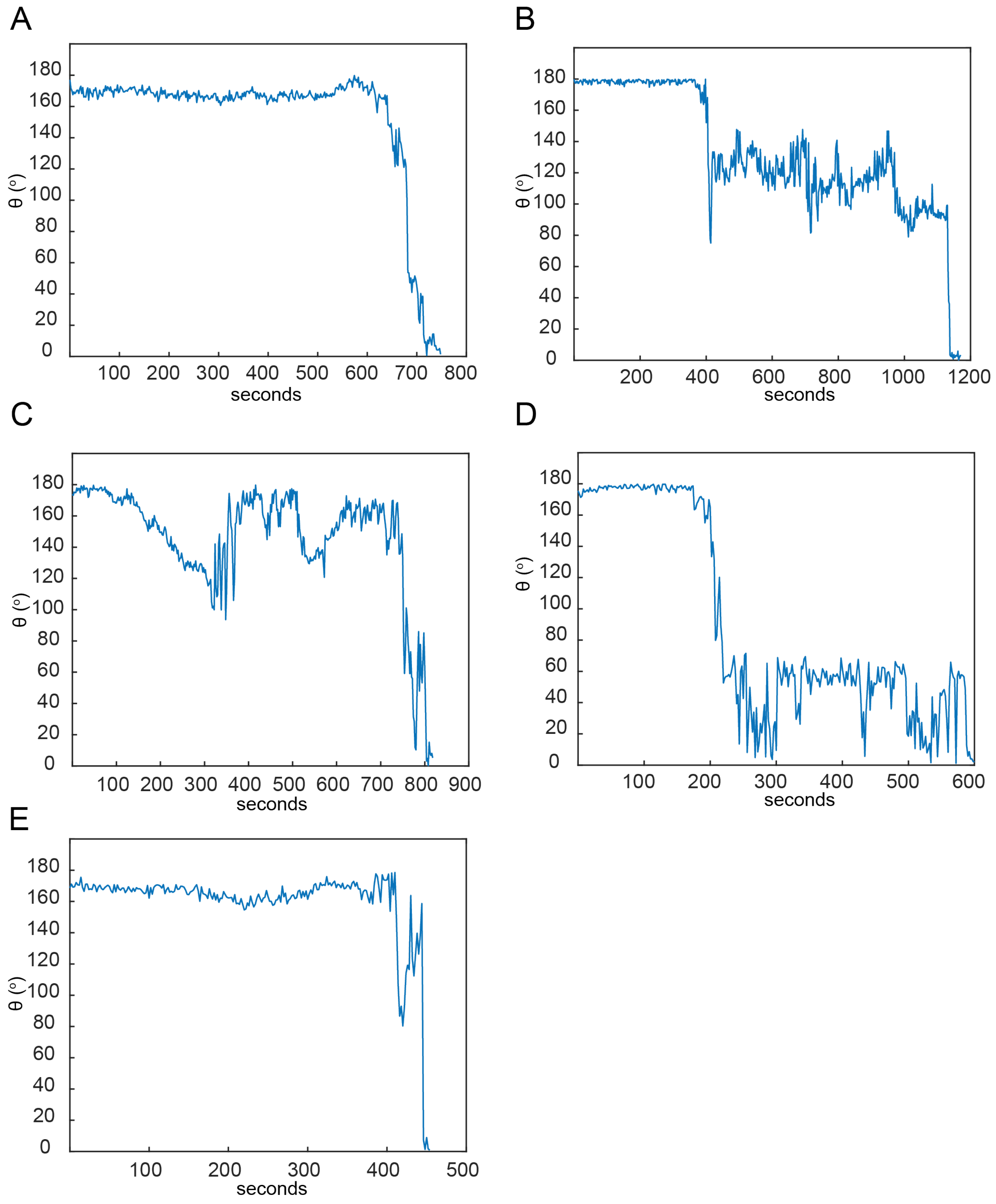


**Figure S17.** Individual examples of sister-cell angle (θ) dynamics during cellular folding (2-second resolution).

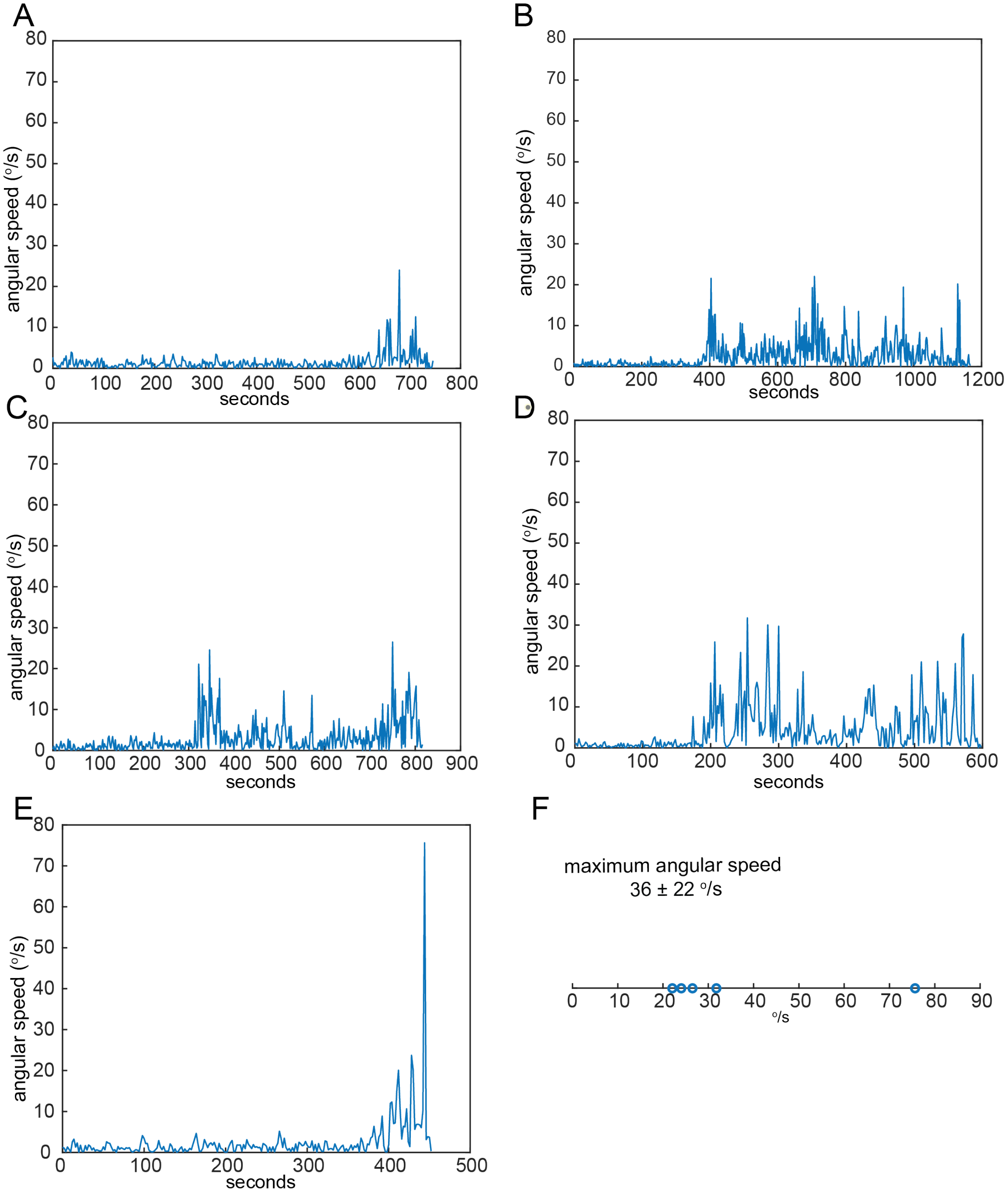


**Figure S18.** (A-E) Individual examples of sister-cell angular speed during cellular folding (2-second resolution). (F) Maximum cell angular speed calculated using data captured at 2-second resolution.

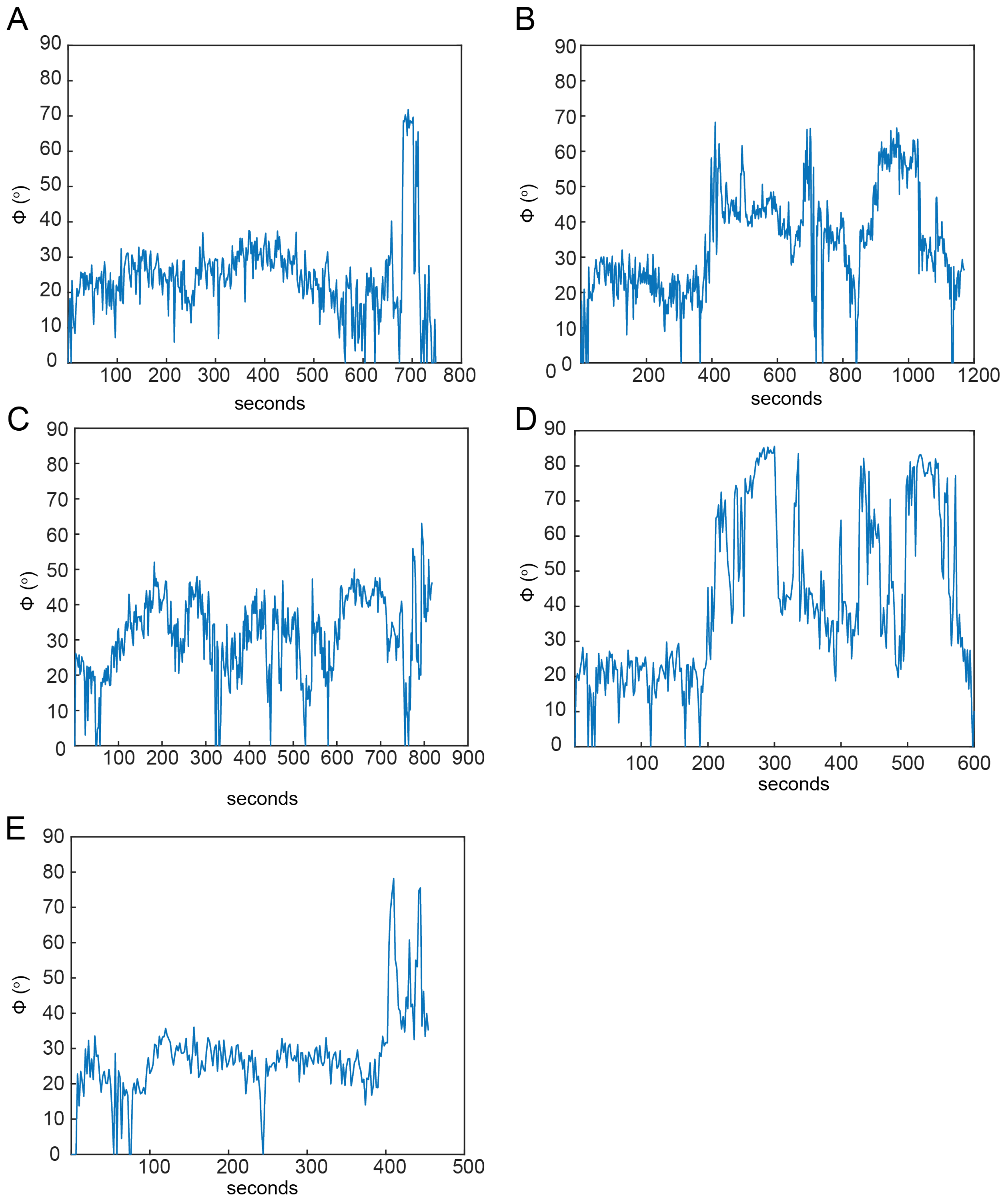


**Figure S19.** Individual examples of cell-surface angle (ϕ) during cellular folding (2-second resolution).

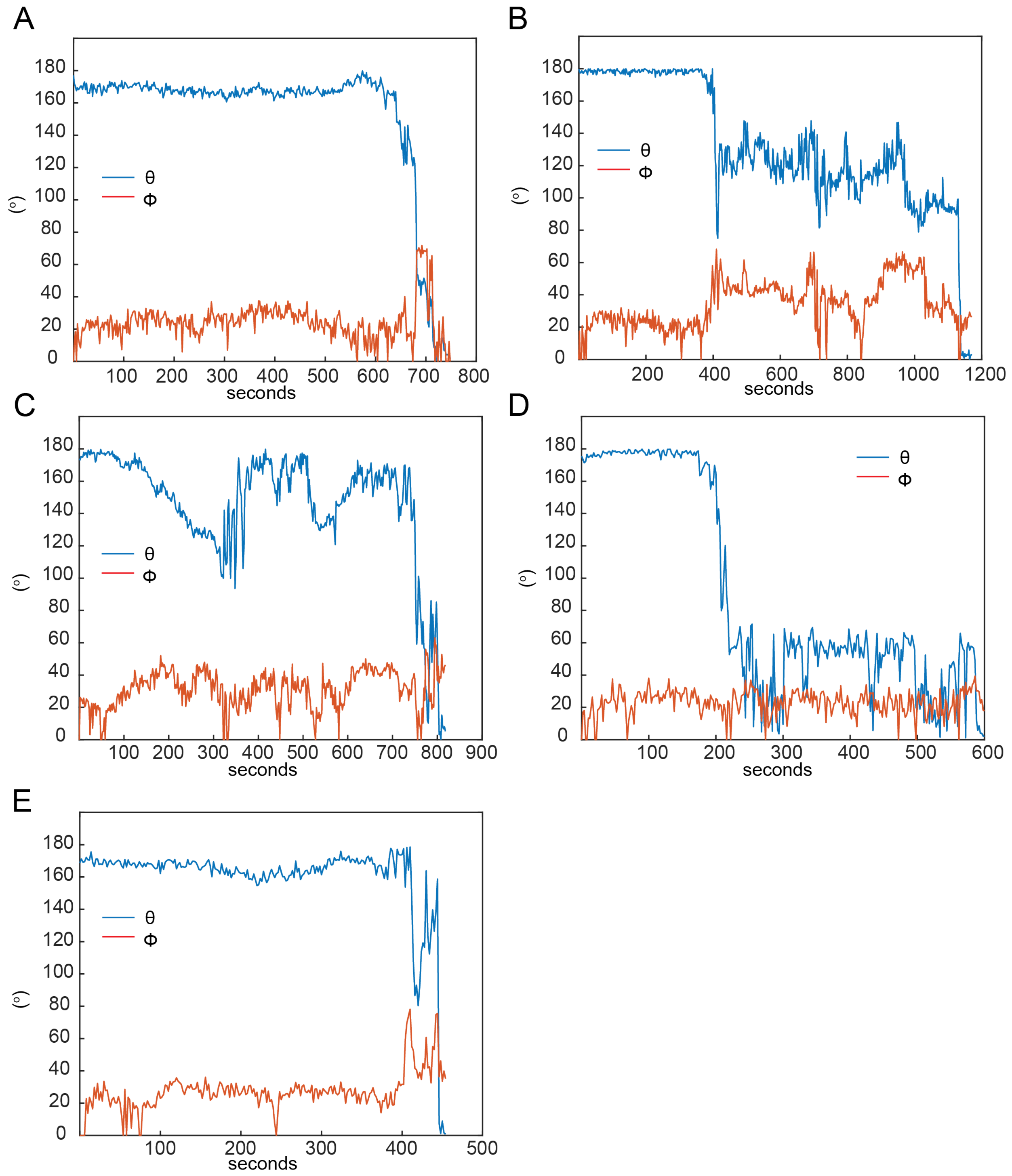


**Figure S20.** Individual examples for dynamics of θ and ϕ (2-second resolution).

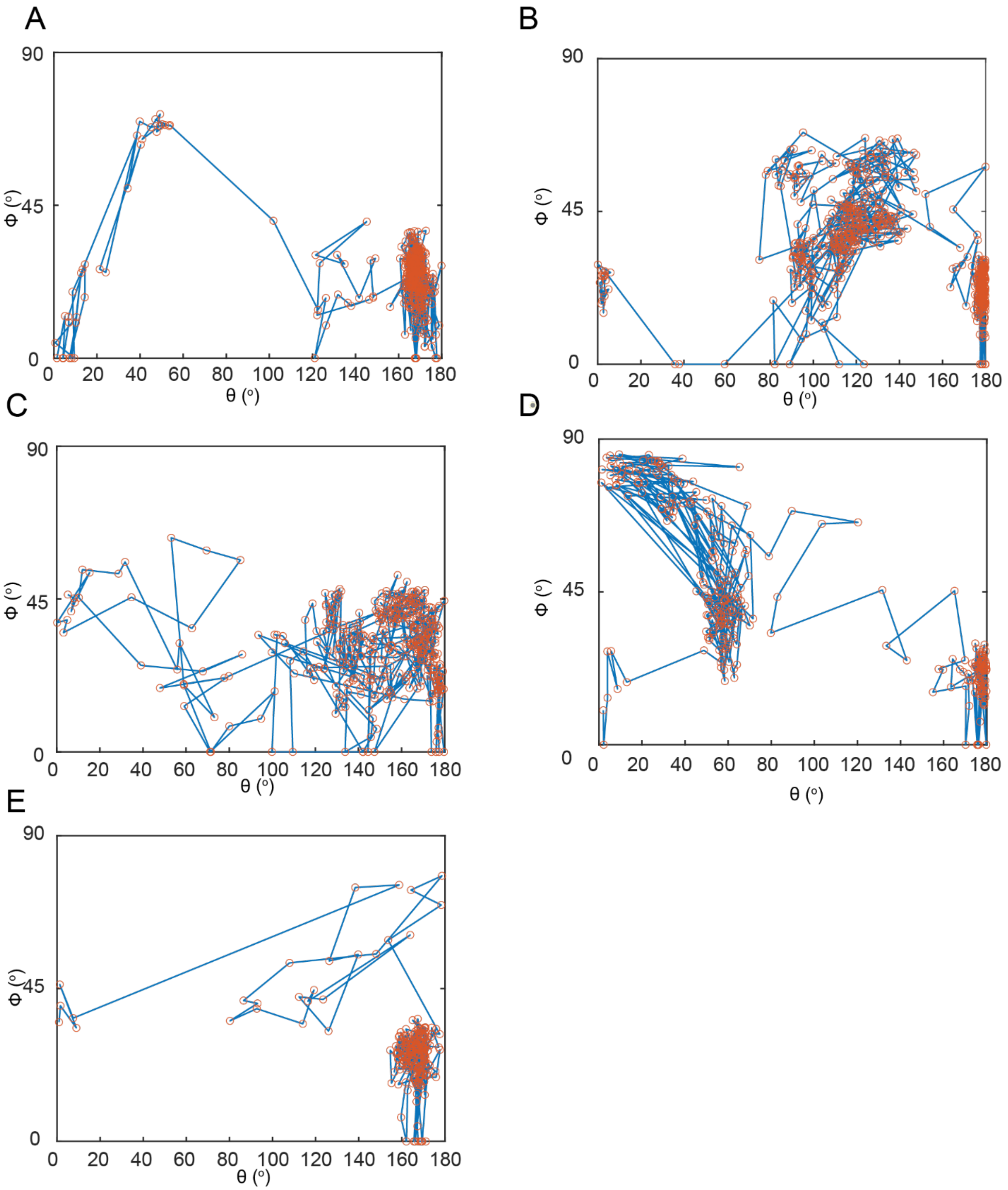


**Figure S21.** Individual examples of cellular folding trajectories in angular coordinates (2-second resolution).

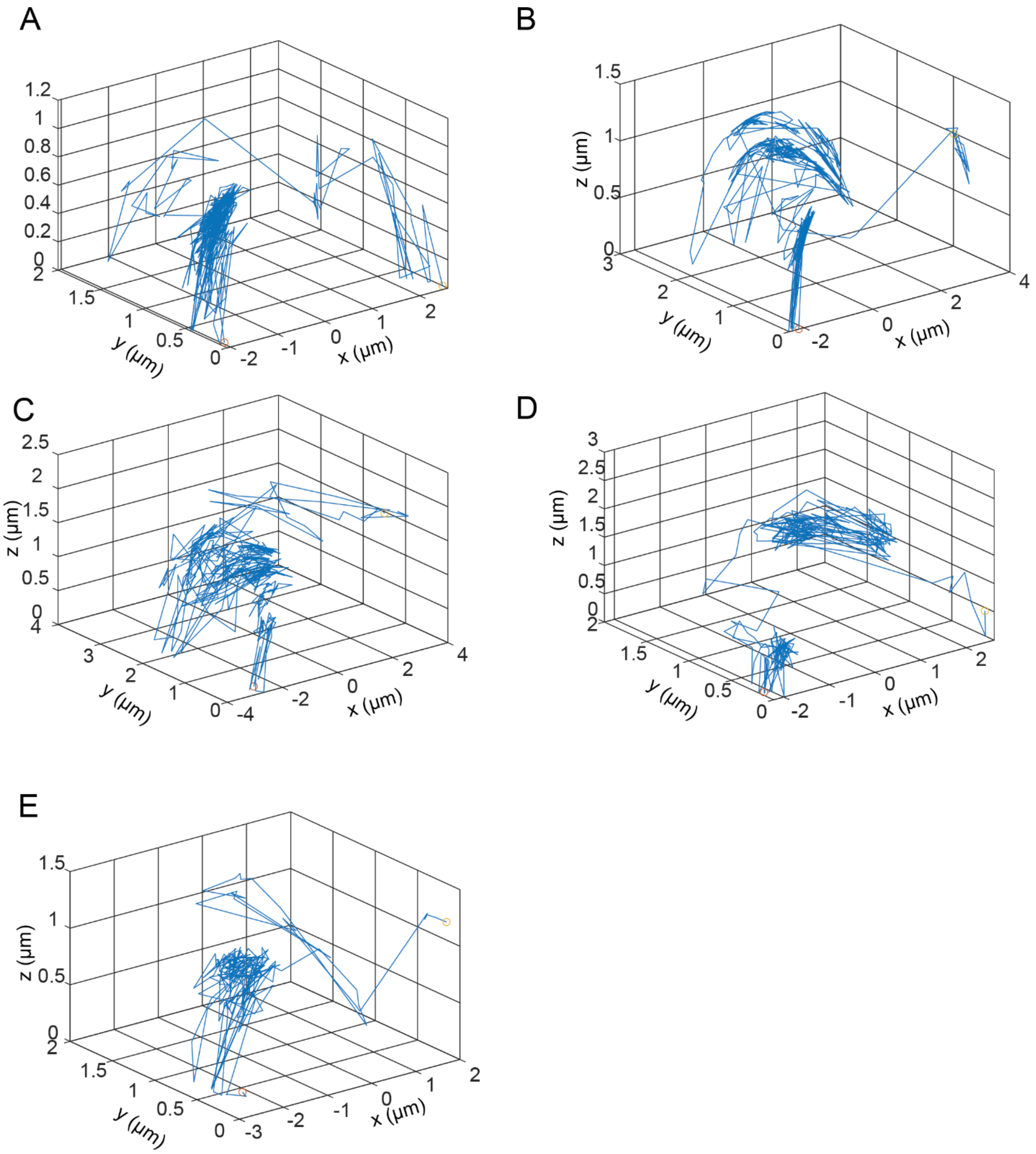


**Figure S22.** Individual examples of cellular folding trajectories in 3D Cartesian coordinates (2-second resolution).

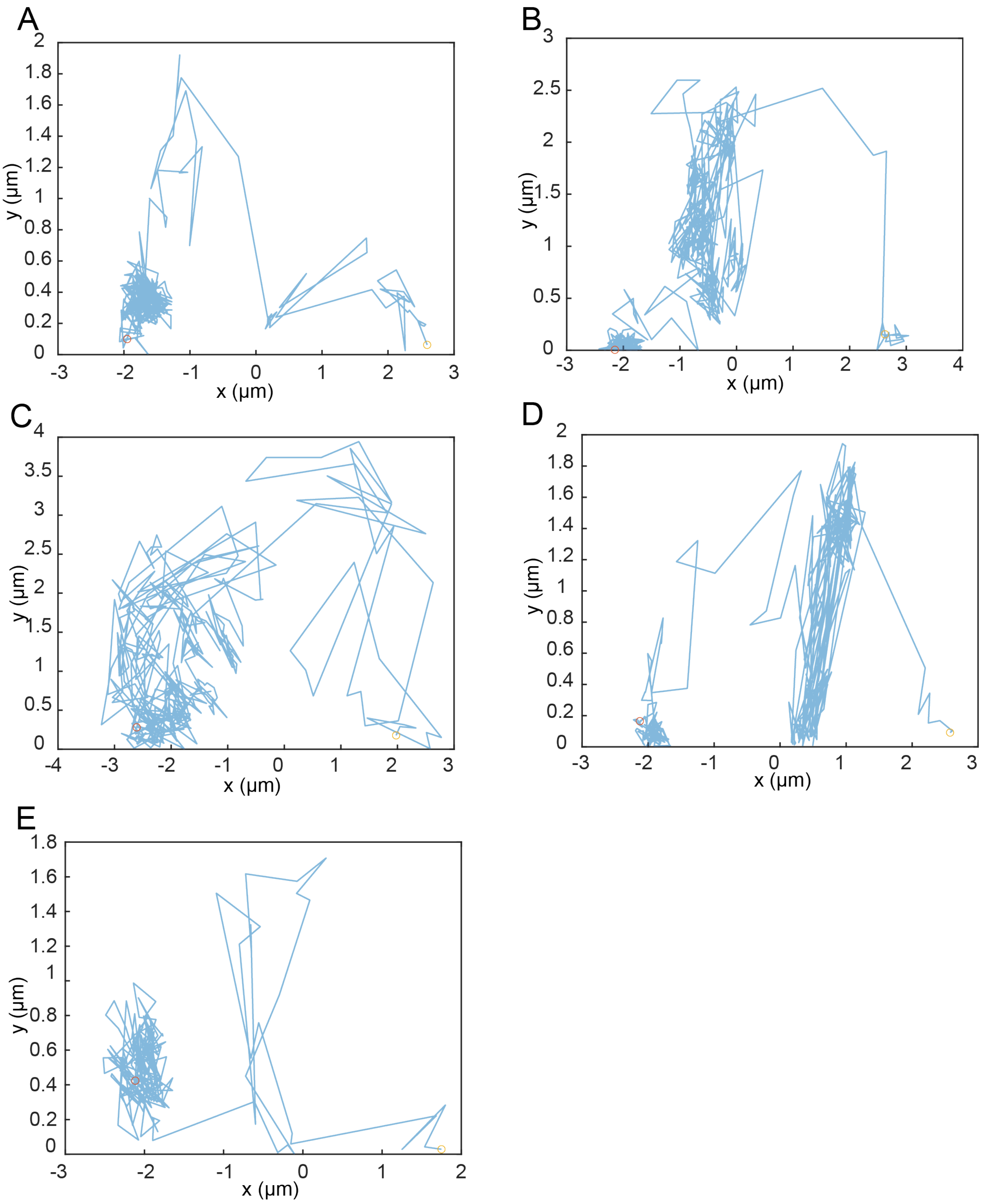


**Figure S23.** Individual examples of cellular folding trajectories in 2D Cartesian coordinates (2-second resolution).

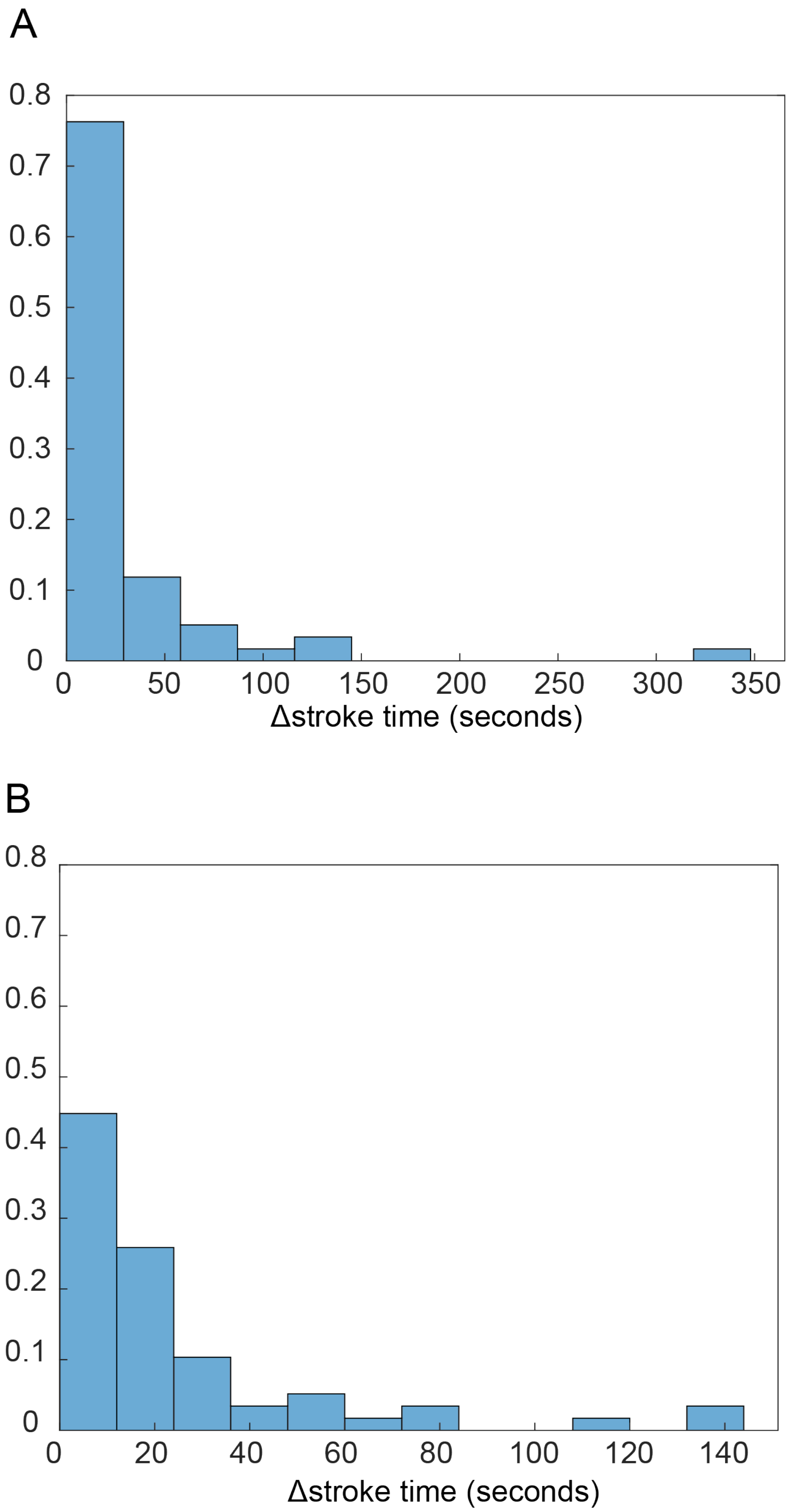


**Figure S24.** Distribution of duration between strokes (A) full distribution and (B) distribution without outliers, *i.e.* durations greater than 250 seconds.

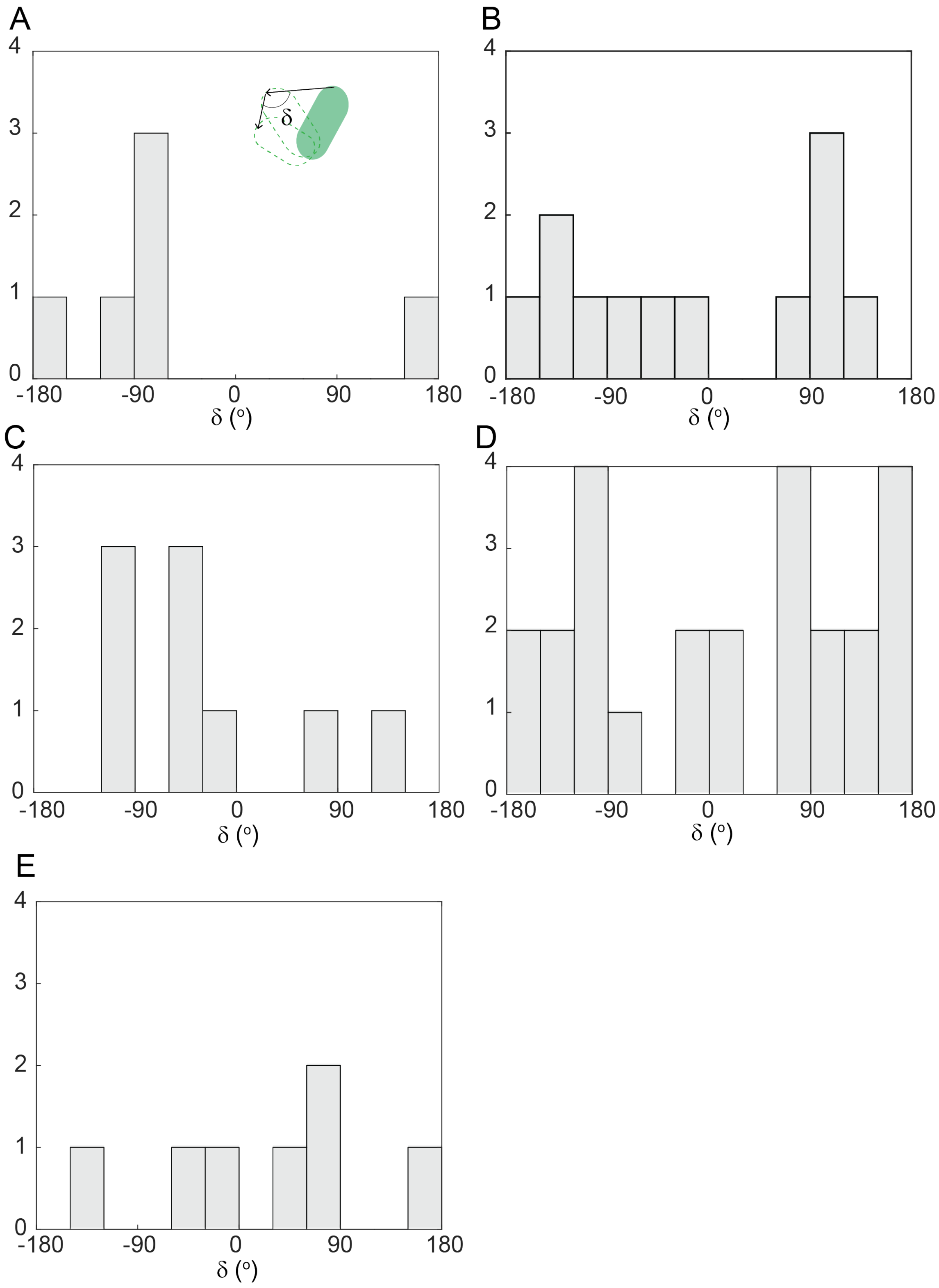


**Figure S25.** Example distributions of angular change between successive strokes for individual cell trajectories. Cumulative distribution plot is indicated in Fig. 1I.

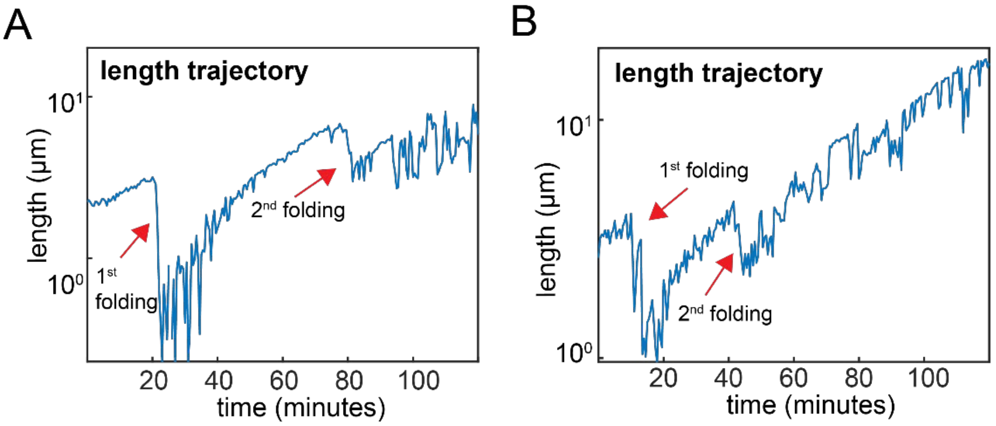


**Figure S26.** Examples of individual community length dynamics, starting from single wild-type cells. Cellular folding is evident when length halves.

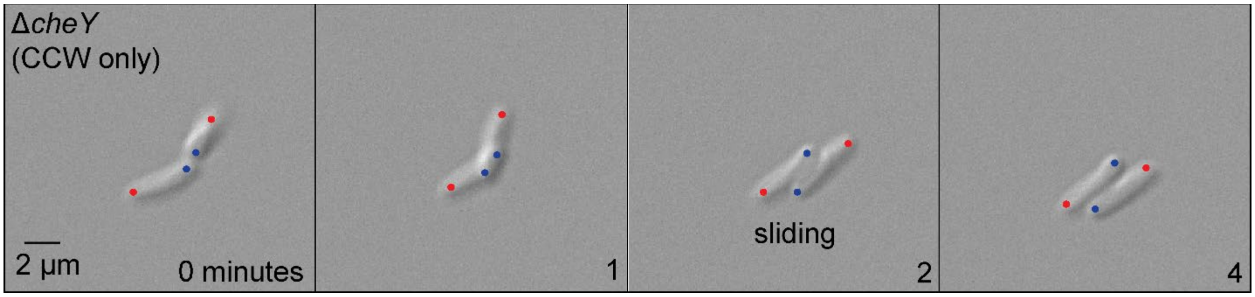


**Figure S27.** Micrographs from representative Movies (2-second resolution) Δ*cheY* cells showing no angular motion and cellular sliding. Time and scale are indicated, as are new (blue) and old (red) poles.

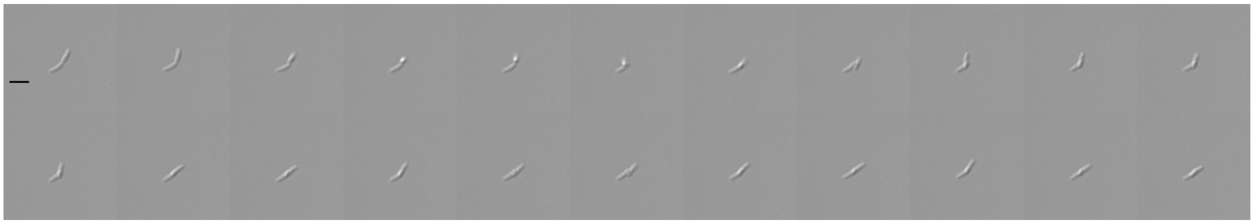

**Figure S28.** Example of *E. coli* Δ*cheY* cell behavior at in DIC at 37°C. Each micrograph is 2 seconds apart and scale bar on first micrograph represents 2 μm.

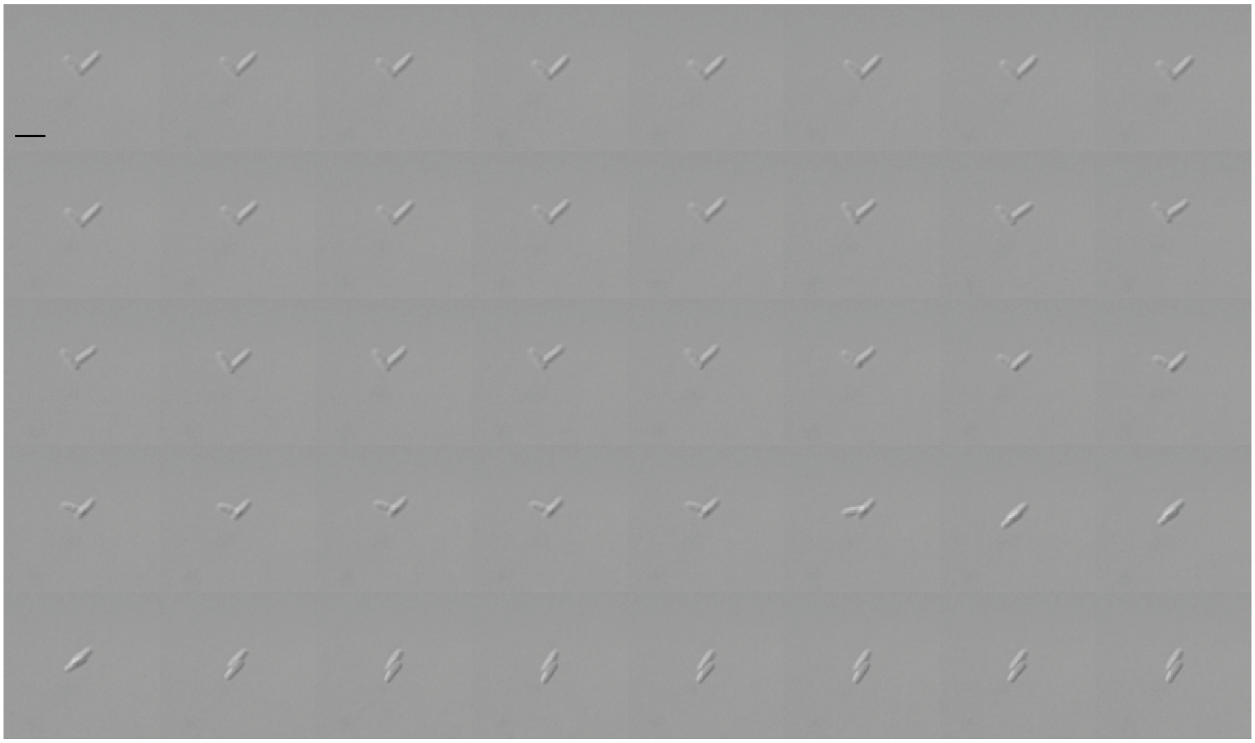

**Figure S29.** Example of *E. coli* Δ*cheY* cell behavior at in DIC at 37°C. Each micrograph is 2 seconds apart and scale bar on first micrograph represents 2 μm.

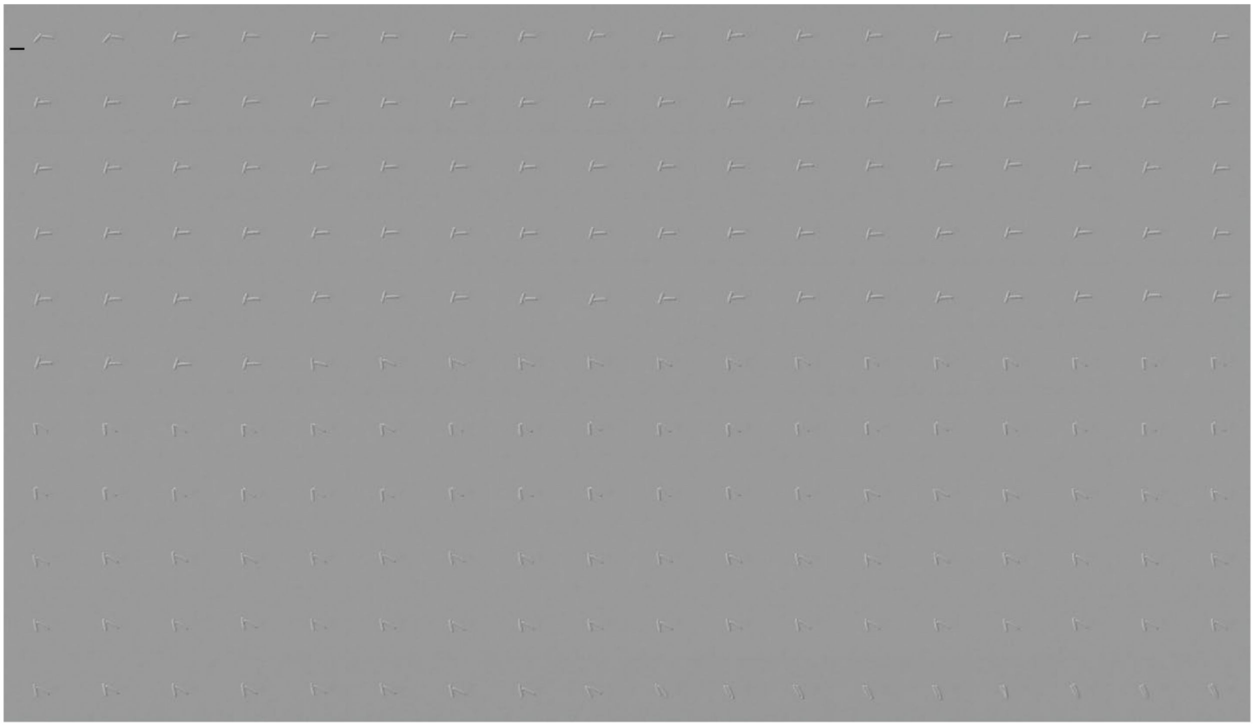

**Figure S30.** Example of *E. coli* Δ*cheY* cell behavior at in DIC at 37°C. Each micrograph is 2 seconds apart and scale bar on first micrograph represents 2 μm.

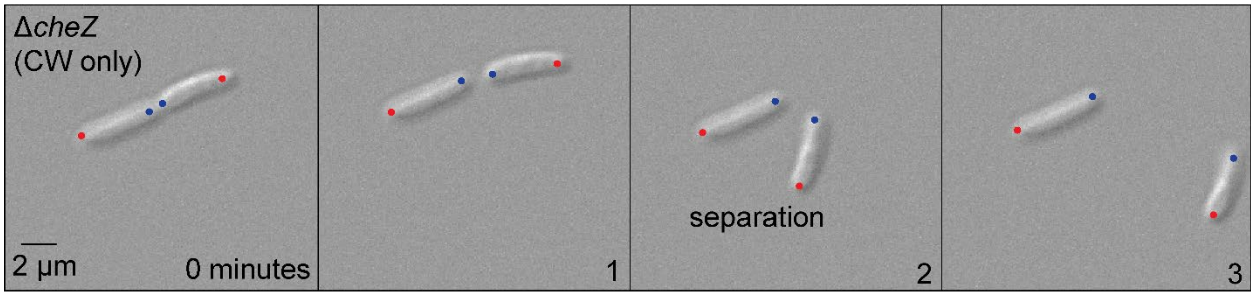


**Figure S31.** Micrographs from representative Movies (2-second resolution) of Δ*cheZ* cells showing significant angular motion and sister-cell dissociation. Time and scale are indicated, as are new (blue) and old (red) poles.

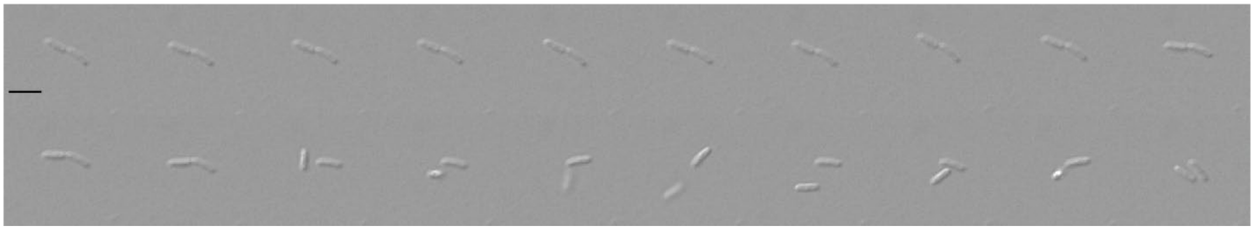

**Figure S32.** Example of *E. coli* Δ*cheZ* cell behavior at in DIC at 37°C. Each micrograph is 2 seconds apart and scale bar on first micrograph represents 2 μm.

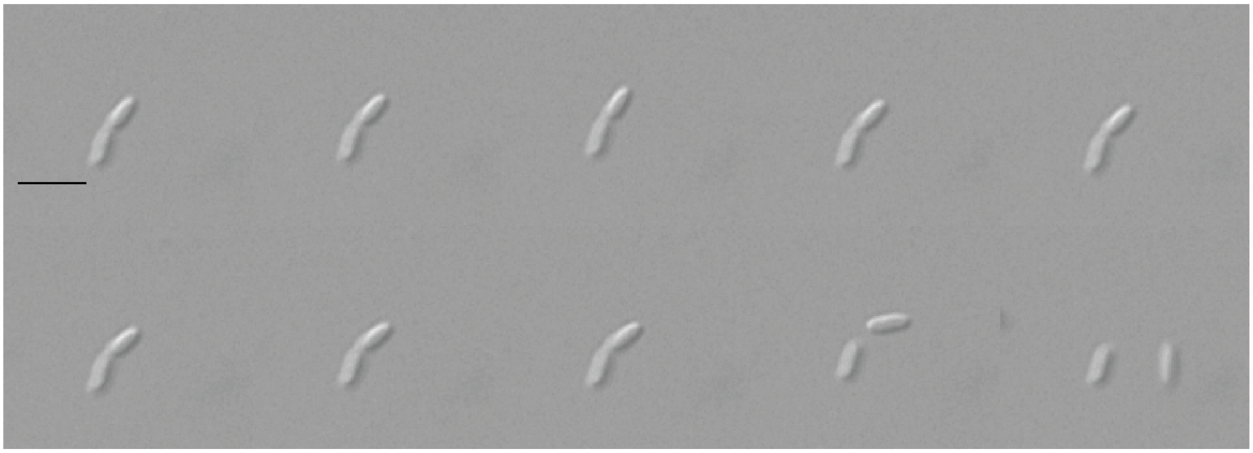

**Figure S33.** Example of *E. coli* Δ*cheZ* cell behavior at in DIC at 37°C. Each micrograph is 2 seconds apart and scale bar on first micrograph represents 2 μm.

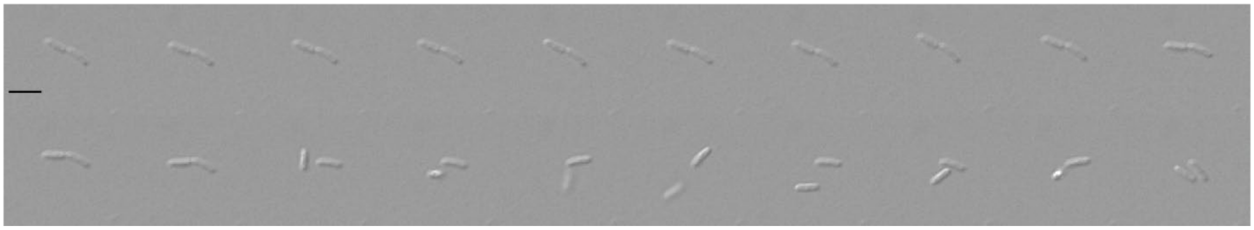

**Figure S34.** Example of *E. coli* Δ*cheZ* cell behavior at in DIC at 37°C. Each micrograph is 2 seconds apart and scale bar on first micrograph represents 2 μm.

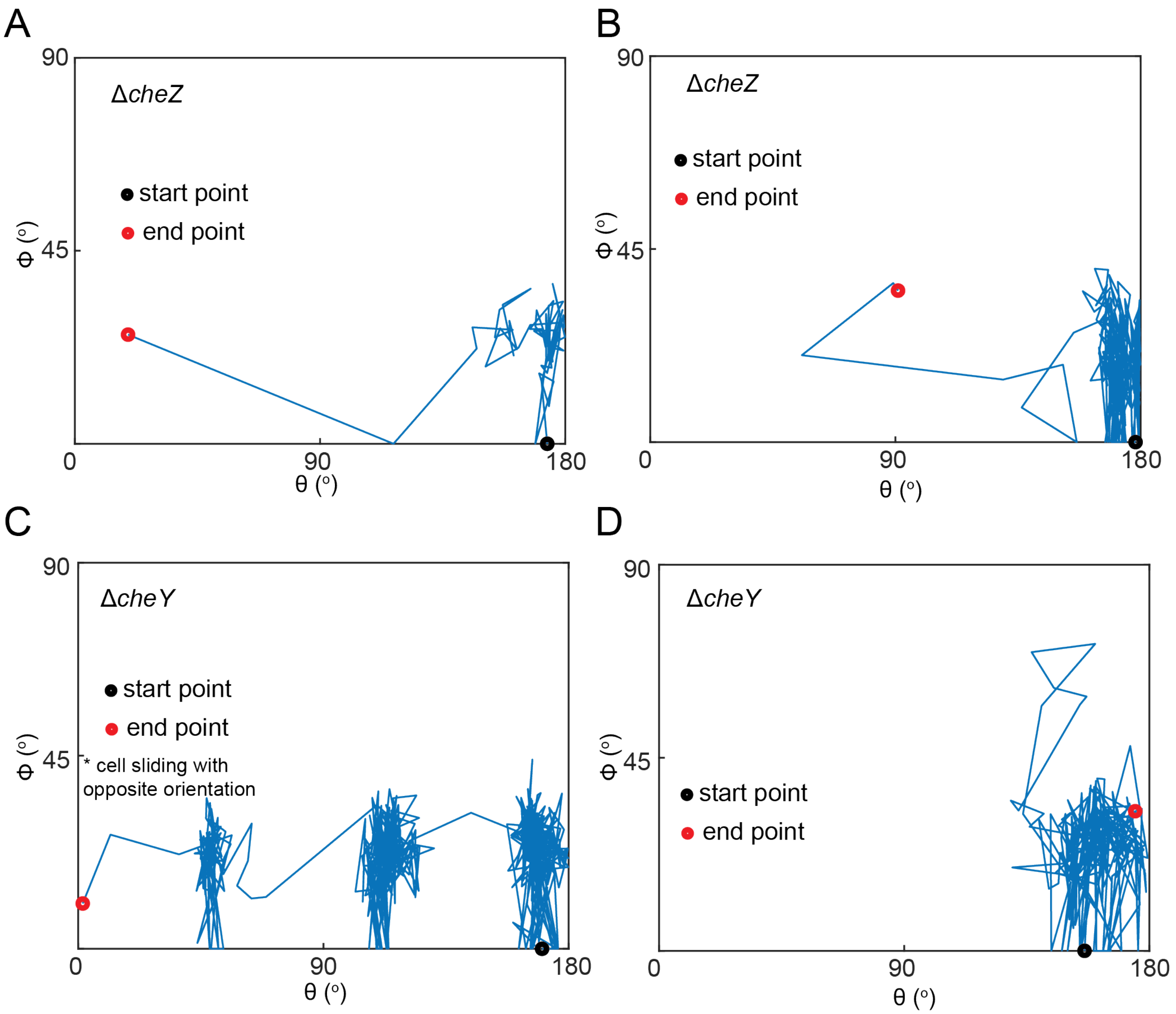


**Figure S35.** Individual examples of cellular motion in angular coordinates for Δ*cheZ* (A-B) and Δ*cheY* (C-D) cells (2-second resolution).

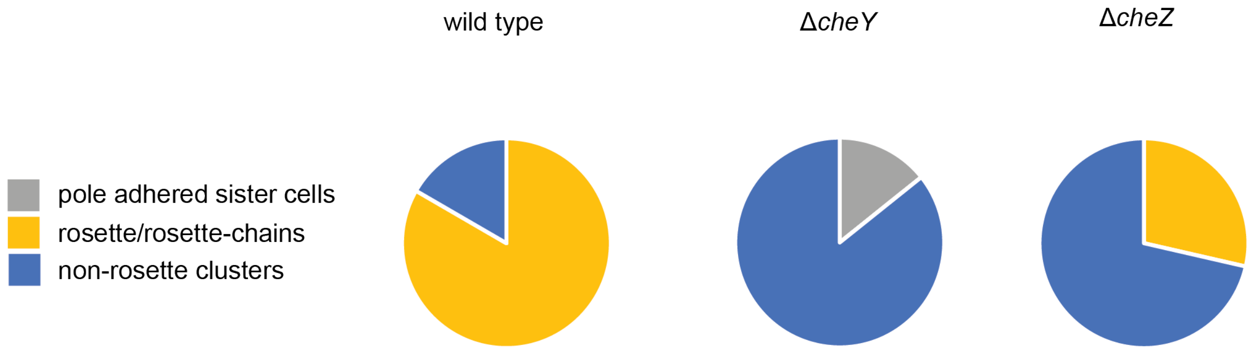


**Figure S36.** Distribution of behavioral fates at 104 minutes for wild-type (n=6), Δ*cheY* (n=7), and Δ*cheZ* (n=7) communities.

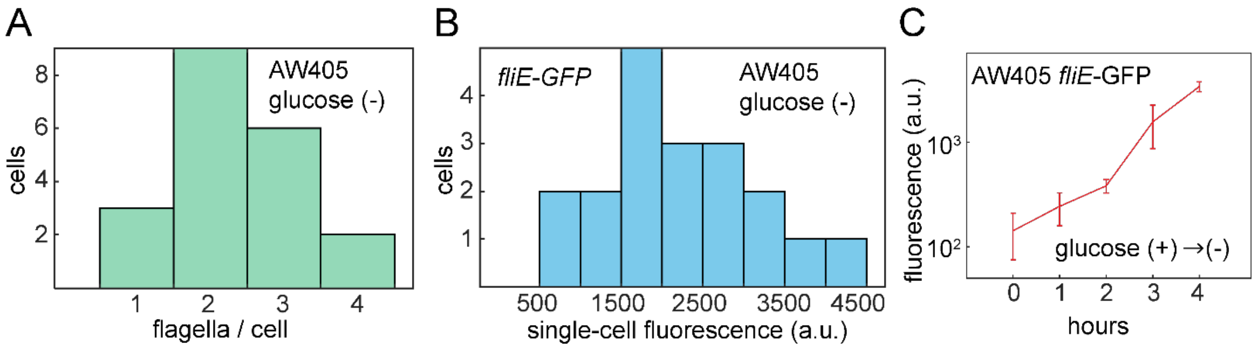


**Figure S37.** **(**B) Distribution of flagella number per cell for AW405 grown in glucose (-) (n=20). Single-cell fluorescence of *fliE*-GFP promoter reporter in AW405 grown in glucose (-) (n=20). (C) Single-cell expression dynamics of *fliE* during glucose (+) to glucose (-) media transition (mean ± standard deviation, n=6). All experiments were performed at 37˚C and 300 RPM before fluorescently imaging individual cells by microscopy.

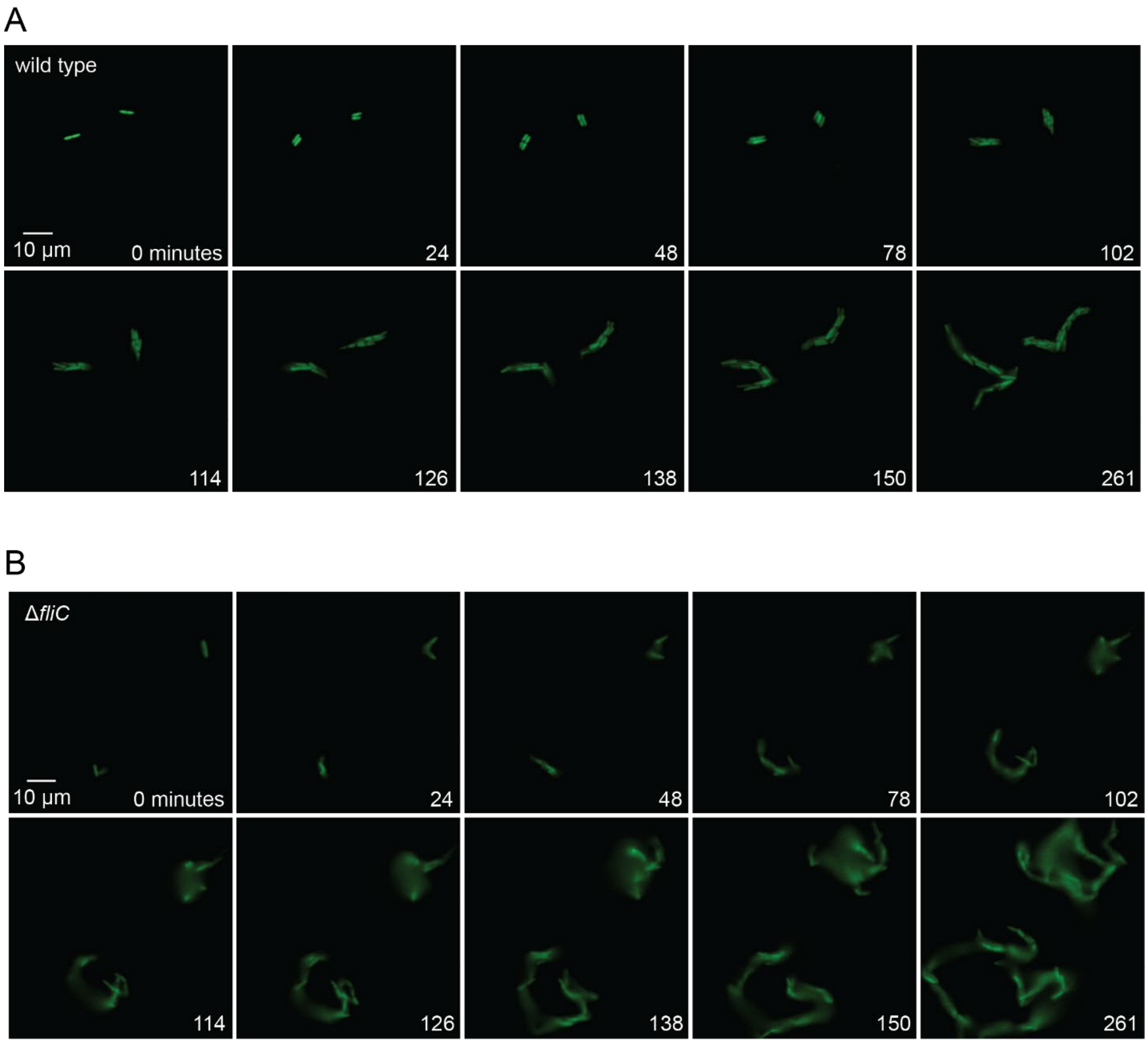


**Figure S38.** (A) Micrographs from example videos of multicellular morphogenesis in wild-type *E. coli* containing GFP. (B) Micrographs from example videos of multicellular morphogenesis in Δ*fliC* *E. coli* containing GFP. Time and scale indicated on micrographs.

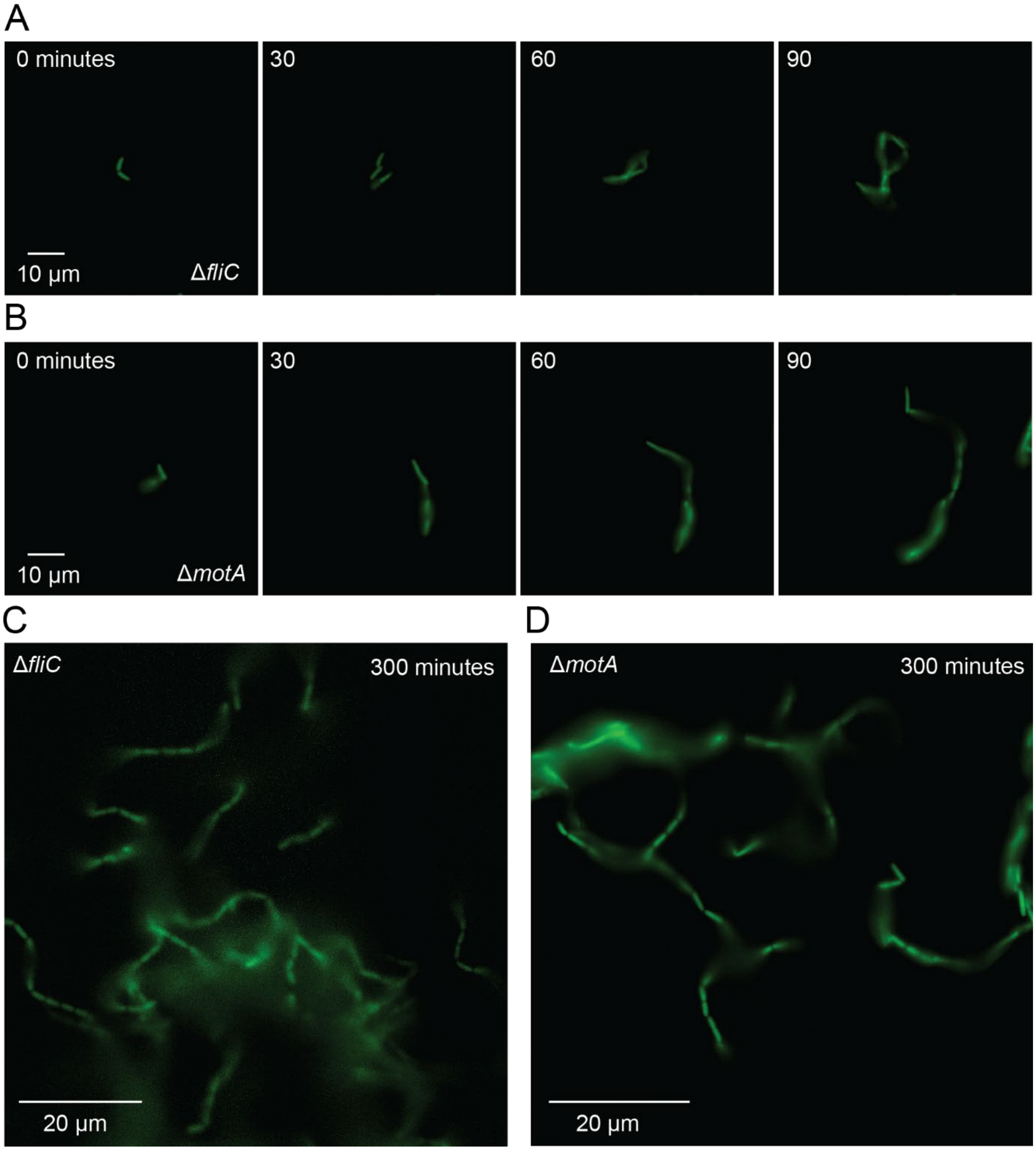


**Figure S39.** Micrographs from example videos of multicellular morphogenesis in Δ*fliC* (A) or Δ*motA* (B) *E. coli* containing GFP. Loose surface attachment in Δ*fliC* (C) or Δ*motA* (D) *E. coli*. Time and scale indicated on micrographs.

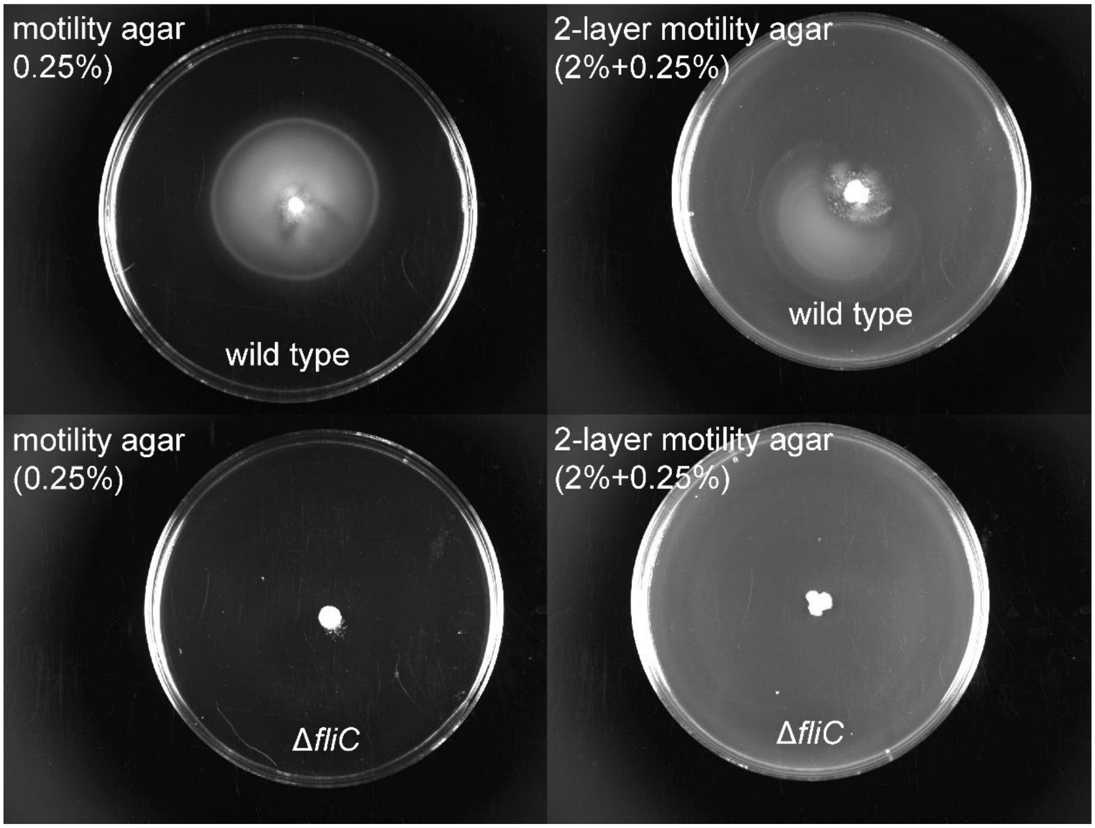


**Figure S40.** Images of bulk-scale motility of wild type and Δ*fliC* cells incubated for 18 hours at 37˚C. Motility agar was made from 20 mL of LB medium with 0.25% agar. 2-layer motility agar was made from a 12 mL bottom layer of LB medium with 2% agar and top layer of 8 mL top layer of LB medium with 0.25% agar.

### Legends of Movies

**Movie S1:** Rosette formation in *E. coli* constitutively expressing GFP. Imaged at 30-second resolution.

**Movie S2:** Multiple examples in single experiment of rosette formation in *E. coli* constitutively expressing GFP. Imaged at 30-second resolution.

**Movie S3:** Cell folding in *E. coli* constitutively expressing green fluorescent protein. Imaged at 30-second resolution.

**Movie S4:** Cell folding in *E. coli* constitutively expressing green fluorescent protein. Imaged at 30-second resolution.

**Movie S5:** Cell folding in *E. coli* constitutively expressing green fluorescent protein. Imaged at 30-second resolution.

**Movie S6:** Cell folding in *E. coli* constitutively expressing green fluorescent protein. Imaged at 30-second resolution.

**Movie S7:** Cell folding in *E. coli* constitutively expressing green fluorescent protein. Imaged at 30-second resolution.

**Movie S8:** Cell folding in *E. coli*. Imaged at 2-second resolution.

**Movie S9:** Cell folding in *E. coli*. Imaged at 2-second resolution.

**Movie S10:** Cell folding in *E. coli*. Imaged at 2-second resolution.

**Movie S11:** Cell folding in *E. coli*. Imaged at 2-second resolution.

**Movie S12:** Cell folding in *E. coli*. Imaged at 2-second resolution.

**Movie S13:** Example of *E. coli* Δ*fliC* cell behavior. Imaged at 2-second resolution.

**Movie S14:** Example of *E. coli* Δ*motA* cell behavior. Imaged at 2-second resolution.

**Movie S15:** Example of *E. coli* Δ*cheY* cell behavior. Imaged at 2-second resolution.

**Movie S16:** Example of *E. coli* Δ*cheY* cell behavior. Imaged at 2-second resolution.

**Movie S17:** Example of *E. coli* Δ*cheY* cell behavior. Imaged at 2-second resolution.

**Movie S18:** Example of *E. coli* Δ*cheZ* cell behavior. Imaged at 2-second resolution.

**Movie S19:** Example of *E. coli* Δ*cheZ* cell behavior. Imaged at 2-second resolution.

**Movie S20:** Example of *E. coli* Δ*cheZ* cell behavior. Imaged at 2-second resolution.

**Movie S21:** Example of cell behavior in *E. coli* AW405 in glucose (-) medium. Imaged at 90-second resolution.

**Movie S22:** Example of cell behavior in *E. coli* AW405 Δ*fliC* in glucose (-) medium. Imaged at 3-minute resolution.

**Movie S23:** Example of cell behavior in *E. coli* AW405 after glucose (+) to (-) media transition. Imaged at 5-minute resolution.

**Movie S24:** Multicellular morphogenesis in wild-type *E. coli* (left) and Δ*fliC* cells (right) expressing GFP. Imaged at 6-minute resolution.

**Movie S25:** Example of wild-type *E. coli* communities on motility agar (0.25% agar) after ~2 hours of growth, imaged at 0.25-second resolution.

**Movie S26:** Additional example of wild-type *E. coli* communities on motility agar (0.25% agar) after ~2 hours of growth, imaged at 0.25-second resolution.

**Movie S27:** Example of wild-type *E. coli* pre-rosette folding with swimming cell on motility agar (0.25% agar) after ~2 hours of growth, imaged at 0.25-second resolution.

**Movie S28:** Example of wild-type *E. coli* cell folding on motility agar (0.25% agar) after ~2 hours of growth, imaged at 0.25-second resolution.

**Movie S29:** Additional example of wild-type *E. coli* cell folding on motility agar (0.25% agar) after ~2 hours of growth, imaged at 0.25-second resolution.

**Movie S30:** Example of Δ*fliC E. coli* communities on motility agar (0.25% agar) after ~2 hours of growth, imaged at 0.25-second resolution.

**Movie S31:** Example of Δ*flu E. coli* cells on motility agar (0.25% agar) after ~2 hours of growth, imaged at 0.25-second resolution.
